# Microglia modulate concussion biomarkers and cognitive recovery in male mice

**DOI:** 10.1101/2025.05.01.651070

**Authors:** Erin D. Anderson, Daunel V. Augustin, Anastasia P. Georges, Taehwan Kim, David A. Issadore, David F. Meaney

## Abstract

There is a critical unmet need for concussion biomarkers that predict cognitive recovery. Existing TBI biomarkers largely capture acute cellular damage rather than the multicellular repair processes that determine long-term outcome, and whether microglia causally shape divergent trajectories remains unclear. Here, we used PLX5622 to deplete microglia and other CSF1R-dependent myeloid cells in a male mouse model comparing concussion alone to concussion preconditioned by prior subconcussive impacts. Microglial depletion had no effect on cognitive outcome after concussion alone, but introduced a significant novel object recognition deficit specifically when concussion was preconditioned, revealing a history-dependent role for microglia in recovery. To identify the molecular substrates of this divergence, we profiled brain-derived GluA1/2+ extracellular vesicle (EV) miRNAs and single-nuclei transcriptomes from the injured brain. Diagnostic and injury-history EV miRNA biomarkers discriminated between concussion subtypes in microglia-intact animals but lost this discriminative power after depletion, indicating that these neuron-enriched biomarkers are influenced by microglia-dependent biology rather than injury alone. snRNA-seq further revealed that microglial depletion selectively induced transcriptional programs related to oligodendrocyte maturation after unconditioned concussion, a candidate substrate for this microglia-dependence. Finally, we identified a serum EV miRNA panel that estimated cognitive recovery across injury conditions. Together, these findings identify a history-dependent role for microglia in concussion outcome and characterize brain-derived EVs as a potentially promising tool for surveilling the neuroinflammatory processes influencing recovery.

## Introduction

The prevailing view in traumatic brain injury (TBI) is that repetitive head impacts are harmful, and for the hundreds to thousands of subconcussive exposures sustained over a contact sports season, this conclusion is broadly supported [1]. Yet evidence from other disorders show that the relationship between acute exposure and outcome is more complex than a simple linear dose-response. For example, preconditioning phenomena use a mild stimulus to confer protection against a subsequent severe insult in preclinical models of ischemic stroke [2], and can involve microglia-dependent mechanisms [3]. We recently demonstrated an analogous effect in TBI: delivering a small number of subconcussive impacts immediately prior to a concussive impact – subconcussive preconditioning – eliminated concussion-associated cognitive deficits in novel object recognition [4]. Preconditioning was time-dependent, protecting only when subconcussive impacts were delivered within minutes of concussion, and was accompanied by a suppression of microglial hyper-ramification and an upregulation of microglial genes inhibiting pro-inflammatory responses, both in the acute phase [4]. These findings suggest a possible causal role for the acute microglial response in mediating cognitive outcomes post-TBI.

Microglia, the brain’s resident immune cells, are increasingly recognized as key mediators of neurological disease progression and recovery [5]. Following injury, microglia clear debris, coordinate a multicellular injury response, and support tissue repair; however, persistent or dysregulated microglial activation creates a neurotoxic inflammatory environment that impedes recovery [6]. As a result, depleting microglia is not necessarily universally beneficial. For example, the outcome of microglial depletion in ischemic stroke depends on its timing, with acute depletion worsening outcomes and delayed depletion improving outcomes [7]. In contrast, in TBI, near-complete microglia depletion initiated prior to injury and throughout the injury and recovery period generally improves cognitive outcomes post-injury [8], including in a model of repetitive mild TBI [9]. Broadly, these effects differ from some effects noted in ischemia models.

Taken together, these observations suggest that the inflammatory state of the brain at the time of concussion could affect cognitive outcome: two patients with the same concussive injury could recover very differently depending on the microglial phenotypes present at the time of injury. However, causal evidence that microglia drive these divergent outcomes is lacking. Moreover, even if this causal relationship were established, we cannot precisely characterize the brain’s inflammatory state from a blood sample using conventional clinical markers, and, critically, use these conventional markers to predict whether that state will lead to successful or impaired recovery.

Existing blood-based biomarkers only partly address this need. Validated blood-based protein biomarkers for mild TBI, GFAP and UCH-L1, predict abnormal neuroimaging within 24-48 hours post-injury [10], whereas other protein biomarkers, such as NfL and S100B, have traditionally been associated with prognosis in moderate-severe TBI [11]. Although S100B has limited utility for mild TBI, recent studies demonstrate NfL has prognostic value for long-term recovery in mild TBI [11–14]. One limitation of these existing biomarkers is that they generally measure endpoints of cellular damage or reactivity, and they do not reveal the biological repair and recovery processes which led to these degenerative endpoints. For this reason, these biomarkers are unable to connect possible damage mechanisms to long-term outcome [15]. Circulating cytokines, perhaps the most direct measure of inflammation, are limited by brief detection windows and poor brain specificity: cytokines originate from many cell types throughout the body [16], and their serum levels typically peak within hours of mild injury and decline rapidly [17], leaving little signal in the post-acute phase when outcome trajectories diverge [18,19]. Circulating miRNAs offer greater mechanistic specificity, with demonstrated potential for diagnostics [20,21], correlation with outcome [22], and persistence of differences months post-injury [23], but lack consensus across studies owing to heterogeneity in injuries and responses [11,24], uncertain cellular origins [25], and sensitivity to confounding factors [11,20,26]. Meanwhile, consensus miRNAs such as miR-21 [20,27] and let-7 [20,28] are nonspecific to TBI or the brain and are broadly associated with disease [29–31].

Cell type-specific biomarkers offer a potentially more precise alternative. Brain-derived extracellular vesicles (EVs) provide a molecular snapshot of their parent cell’s state and play a key role in intercellular signaling [32]. Across neurological diseases, neuron-enriched EVs contain cargo indicative of disease severity [33,34], cognitive impairment [35,36], and treatment response [37,38]. For TBI, prior work using GluA1/2+ EVs has classified diagnosis and injury history, although without connecting biomarker signatures to cognitive outcomes [39–41]. Meanwhile, emerging efforts to characterize microglia-enriched EVs in neurological disease have relied on EV abundance rather than content [42,43]. In this study, we use neuron-enriched GluA1/2+ EVs to capture not what microglia are signaling, but how neurons respond to the inflammatory milieu — an integrative readout of the multicellular recovery environment that could tie more directly to cognitive function.

In this work, we consider whether neuron-enriched GluA1/2+ EVs provide insight into neuroinflammation based on concussion subtype and estimate prior cognitive recovery. First, we use GluA1/2+ EV miRNAs to identify diagnostic and injury history miRNAs, using a model for single, unconditioned concussion compared to concussion preconditioned by subconcussive impacts. To evaluate whether microglia-dependent effects are reflected in neuron-enriched blood-accessible biomarker signatures, we use PLX5622 to acutely deplete microglia and other CSF1R-dependent myeloid cells and then compare GluA1/2+ EV miRNA content across injury subtypes. We then use single-nuclei transcriptomics as a complementary metric to characterize some of the persistent cellular responses underlying injury recovery. We test microglia’s involvement in acute cognitive recovery differs based on prior injury history, and, finally, we identify an EV miRNA panel correlated with prior cognitive recovery across injury conditions. Together, these results suggest a context-dependent role for microglia in concussion outcome and identify brain-derived EVs as a potentially valuable tool for surveilling the neuroinflammatory processes that influence recovery.

## Methods

### Animals

In this study, we used 8-12 week old male C57BL/6 mice (Charles River Laboratories). We focused on male mice due to their more rapid and pronounced immune response following traumatic brain injury compared to female mice [44–46]. Moreover, there is evidence that male mice benefit from early microglial attenuation post-injury more than female mice [47]. Including both sexes would have required a fully powered factorial design beyond the scope of this preliminary mechanistic study. Future work will examine whether these findings generalize to females. Mice were housed in a facility with a 12h light/dark cycle with *ad libitum* access to food and water. All protocols were approved by the University of Pennsylvania Institutional Animal Care and Use Committee (IACUC).

### Experimental Design

#### Closed-head Controlled Cortical Impact

To deliver subconcussive and concussive impacts, we used closed-head controlled cortical impact as previously described [4]. Briefly, we initiated anesthesia with 3.5% isoflurane gas in oxygen until the mouse reached a deep plane of anesthesia and then maintained anesthesia at 1.5-2%. Following scalp incision, we used a 6-mm diameter rubber impactor centered over the left parietal bone to deliver either a concussive impact at 3.5 m/s or a subconcussive impact at 1 m/s. Sham animals received no impact. For all impact speeds, the impactor tip was in contact with the skull for 30 ms, achieving a peak impact depth of 1 mm for both impact protocols. We studied two injury paradigms: either a Concussion (C) consisting of a single 3.5 m/s impact, or a Preconditioned Concussion (PC) consisting of two subconcussive (1.5 m/s) impacts delivered followed by a single concussive (3.5 m/s) impact (Fig 1A), consistent with our prior study [4]. Recent work from our group showed that a single concussive impact can cause novel object recognition deficits 4 days after impact, while administering two subconcussive impacts prior to this concussive loading (i.e., preconditioned concussion) would reverse these deficits [4]. This finding, that rapid preconditioning prevented cognitive deficits associated with concussion, led us to consider whether the effect was associated with microglial activity and indeed whether there was a relevant biomarker for the effect. Mice were randomly assigned to each injury condition. Mice were excluded from the study if they exhibited a skull fracture post-impact. Following the final impact in either injury protocol, we sutured the mouse’s scalp, placed them in a heated recovery cage until they were alert and mobile as normal, and then returned them to the colony.

**Fig 1.**
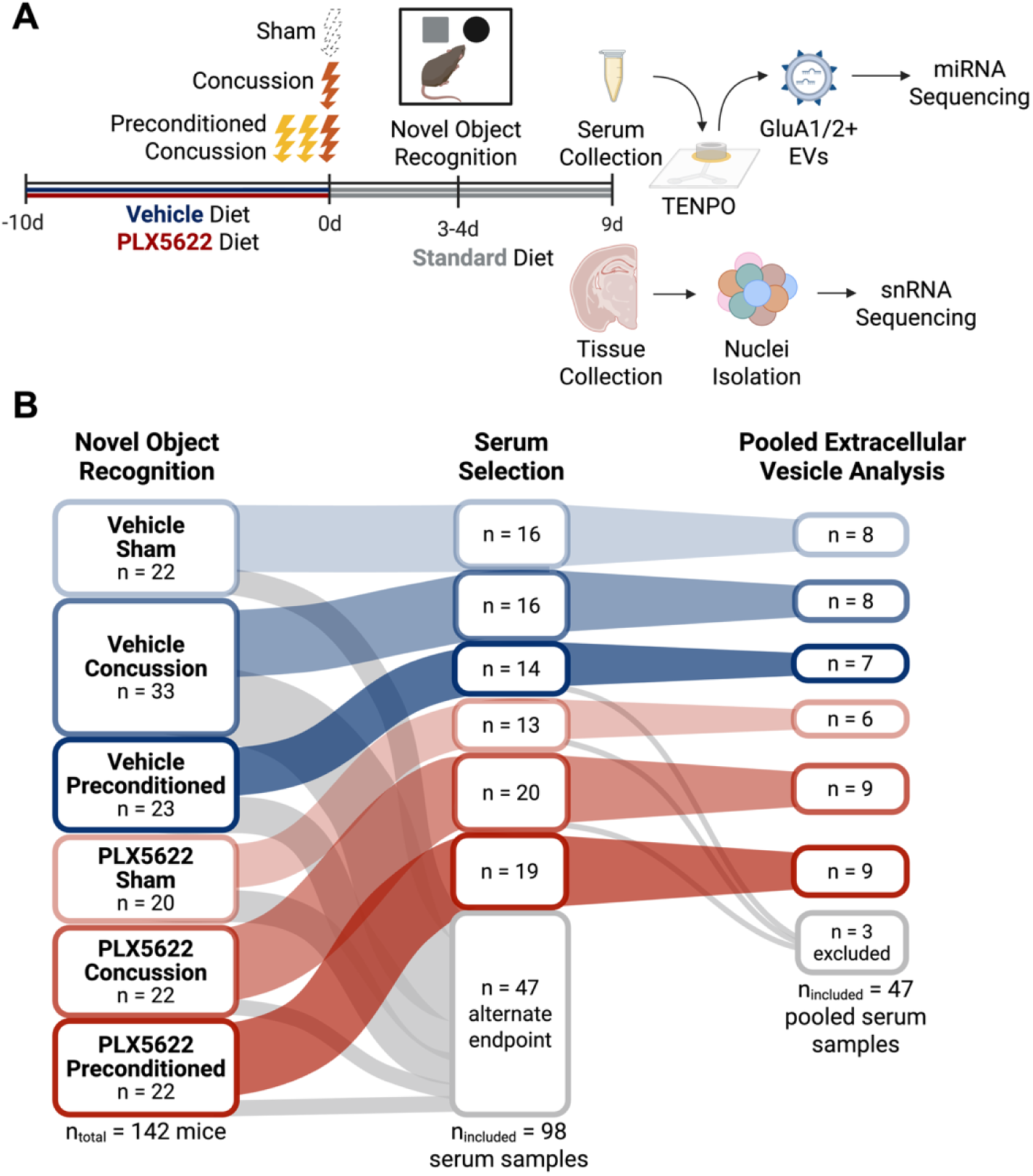
Experimental timeline and sample allocation. (A) Experimental timeline - Vehicle or PLX5622 diet was administered for 10 days leading up to injury. We considered sham, single concussion, and concussion preconditioned by two subconcussive impacts. We conducted Novel Object Recognition on Days 3-4 post-injury and collected serum for extracellular vesicle (EV) analysis or brain tissue for single nuclei RNA-seq at 9 Days post-injury. (B) Flow of samples through study - serum samples derived from the 142 mice included in NOR analysis (N = 20-33/condition) were pooled into 47 total samples for extracellular vesicle (EV) analysis (N = 6-9/condition), leaving 47 samples behind for alternate endpoints (e.g., perfusion). Figure generated using BioRender.

#### PLX5622 Treatment

To deplete microglia, we administered PLX5622 (Amadis Chemical) incorporated into AIN-76A semi-synthetic rodent chow at 1200 mg/kg by Research Diets, consistent with prior studies [48,49]. Mice were assigned on a cage-by-cage basis to *ad libitum* access to either PLX5622-incorporated or vehicle chow for 10d prior to injury (Fig 1A), which was sufficient to deplete microglia, as verified in prior studies [48,50–52]. Each experimental batch contained animals from each diet condition. In preliminary studies, we ensured this dosage and time was sufficient to deplete >90% of microglia (Fig S1). Following injury, we returned mice to the standard grain-based diet, Lab Diet 5001 (Animal Specialties and Provisions).

### Immunohistochemistry

Immunohistochemistry (IHC) was performed as described in [4]. Briefly, we used IHC to confirm microglial depletion (Fig S1). Microglia and macrophages were stained using anti-Iba1 (Wako; 019-19741) in 20 μm coronal brain slices following 10 days of vehicle or PLX5622 treatment.

### Serum collection

On Day 9, we collected serum for extracellular vesicle (EV) isolation (Fig 1A). We selected day 9 post-injury so that cognitive assessment could be conducted prior to the terminal blood draw and EV content could be correlated with prior cognitive recovery (see: Methods: Behavioral Testing). To collect blood, we administered a lethal dose of sodium pentobarbital (250 mg/kg) and then collected whole blood via cardiac puncture. We allowed the whole blood to coagulate in uncoated tubes at room temperature for 30-60 minutes and then centrifuged at 433 x *g* for 15 min at 4C to isolate clear serum. We stored the serum at -80C until processing.

### GluA1/2+ extracellular vesicle isolation from pooled serum

To isolate GluA1/2+ extracellular vesicles (EVs), we used nanofluidic <u>T</u>rack <u>E</u>tched <u>N</u>ano<u>PO</u>re (TENPO) devices as previously described [40,41] with a 3-μm pore size [53]. Briefly, we thawed serum and pooled 1-4 mouse serum samples immediately prior to TENPO processing, excluding serum samples with visible red blood cell contamination. The current TENPO design requires ≥ 500 µL of serum, and the typical serum volume derived from each animal averaged 220 µL. Therefore, we developed a sample pooling process that grouped serum samples with similar novel object preference scores (average range within 6.7 points), completing the pooling when the pooled sample volume was ≥500 µL, and resulting in 50 total serum samples with n = 7-9 pooled samples per experimental condition, consistent with prior work [40]. Pooling using similar frameworks (i.e., within-condition, then further stratified based on behavior) has been used previously in preclinical models of impaired versus unimpaired aging [54] and stress-susceptible versus stress-resilient mice [55]. Although pooling can intrinsically mask outliers, this concern is partially mitigated by only including animals with novel object preferences within a normal distribution (see: Methods: Behavioral Testing) and restricting to clear serum without red blood cell debris that could introduce variability. Across the 47 serum samples included in our analysis (see Methods: Differential miRNA expression, Fig 1B), 2 were from one animal apiece, 41 were from two animals, 3 were from 3 animals, and one was from 4 animals. Pooling ensured consistent performance across TENPO chips and downstream RNA extraction efficiency, allowing us to detect subtle differences in EV miRNA expression. We labeled neuron-enriched EVs by incubating the serum with biotinylated anti-GluA1/2 antibodies (Bioss; bs-10042R-Biotin). GluA1 and GluA2 are AMPA-type glutamate receptor subunits primarily expressed by neurons, but also expressed by other brain cells. Prior work demonstrates that GluA1/2+ EVs are enriched in neuronal media over astrocyte media [56]. We prepared GluA1/2-labeled EVs for magnetic capture on chip by incubating with anti-biotin ultrapure microbeads (Miltenyi-Biotec). We ran the labeled and microbead-conjugated serum samples through the TENPO devices (Chip Diagnostics) using a programmable syringe pump to extract the GluA1/2+ EVs as described in Lin et al., 2023 [53]. We lysed the GluA1/2+ EVs on chip using QIAzol (Qiagen) to extract the RNA content.

### miRNA isolation and sequencing

To isolate miRNAs from the extracellular vesicle lysate, we used the miRNeasy kit (Qiagen) and stored the miRNA at -80C until library preparation. We used the QIAseq miRNA Library Kit (Qiagen) to produce the miRNA library. We sequenced the library using the NovaSeq 6000 SP Kit (Illumina; Next-Generation Sequencing Core, University of Pennsylvania). We demultiplexed the raw sequencing data for each sample using the BCL Convert v2.4 application from Illumina to generate FASTQ files. We aligned the FASTQ files to the *Mus musculus* genome in miRBase_v22 using Qiagen’s GeneGlobe Analysis Portal. We exported the expression matrices for each sample for processing in RStudio.

### Differential miRNA expression

Prior to differential miRNA expression analysis, we excluded 3 pooled serum samples which failed library preparation with a total miRNA read count less than 10, and excluded miRNAs which had an average read count per sample less than one [57]. We then used DESeq2 (Version 1.44.0) for R (Version 4.4.1) to normalize the data and identify differentially expressed miRNAs with an FDR-adjusted Q < 0.1. Due to the exploratory nature of this work, we used Q < 0.1 rather than Q < 0.05 to prioritize sensitivity. Data are presented as log2 of the normalized read counts plus a nominal value (0.5 - DESeq2 default) to ensure a mathematically valid output.

### Evaluation of diagnostic and injury history biomarkers across diet conditions

We identified candidate diagnostic and injury history biomarkers *a priori* by differential expression analysis as described previously (see: Methods: Differential miRNA expression). To determine whether these candidate biomarkers generalized between vehicle and microglia-depleted (PLX5622) conditions, we generated separate logistic regression models for each candidate miRNA using expression as the sole predictor of injury status. We selected logistic regression because it provides an interpretable estimate of the relationship between a single biomarker and a binary outcome while avoiding the additional model complexity of multivariable classifiers. We trained logistic regression models using data from one diet condition (vehicle or PLX5622), and applied the fitted model coefficients (without retraining) to the opposite diet condition to assess whether biomarkers retained their ability to discriminate injury status across dietary conditions. Model discrimination was quantified using the area under the receiver operating characteristic curve (AUC), and 95% confidence intervals were estimated using the DeLong variance estimator. Given the modest sample size, we report confidence intervals for the AUC to characterize the precision of discrimination estimates rather than performing formal statistical comparisons between AUCs, which would have limited power and unstable estimates.

### Nuclei isolation and sequencing

At 9 days post-injury, we administered 250 mg/kg sodium pentobarbital (i.p.) to the mice, quickly removed the brain from the skull, and collected a 1-mm ipsilateral slice of brain tissue directly below the site of injury (-1.5-2 mm from bregma), which was flash-frozen and stored at -80C until nuclei isolation. We isolated nuclei from n = 4 samples/condition, consistent with prior work [49,58,59]. To isolate nuclei, we first homogenized tissue according to 10X Genomics Demonstrated Protocol CG000214 Rev F: Nuclei Isolation from Cell Suspensions and Tissue for Single Cell RNA Sequencing - Mouse Brain Tissue. We next removed myelin using an iodixanol gradient [60]. Finally, we fixed nuclei using the Evercode Nuclei Fixation v3 Kit (Parse Biosciences) and stored nuclei at -80C until library preparation. We prepared the library using the Evercode WT v3 Kit (Parse Biosciences) and sequenced using the NovaSeq 6000 SP Kit (Illumina) at Penn’s Next Generation Sequencing Core.

### Single nuclei RNA sequencing (snRNA-seq) analysis

Data processing and analysis were performed using Trailmaker v1.2.1 (Parse Biosciences) embedded with ScanPy [61]. We used Trailmaker’s automated data processing methods to exclude nuclei with low transcript number and doublets. We annotated nuclei by harmonizing leiden clusters with CellTypist’s automated logistic regression-based annotation using the Allen Mouse Brain Cell Atlas [62,63], omitting one cluster for uncertain cellular identity. Differentially expressed genes are identified using a pseudobulk limma-voom workflow in the Trailmaker software. We accounted for multiple comparisons using false discovery rate (FDR) and report RNAs with FDR < 0.1.

### Functional enrichment analysis

For functional (pathway) enrichment analysis for EV miRNAs, we used miRNet v2.0 for miRNAs matched to target genes based on miRTarBase v9.0 and pathways using Gene Ontology (GO) Analysis, focusing on Biological Processes (BP). We used a hypergeometric test to control for multiple comparisons with a false discovery rate (FDR) < 0.05. We included the top 10 terms with the largest enrichment ratios in the main text and provide complete GO: BP lists in the supplement.

For RNAs identified through snRNA-seq, we employed over-representation analysis (ORA), using EnrichR’s GO Enrichment Analysis separately for upregulated and downregulated gene sets by comparison, applying Fisher’s Exact Test with FDR < 0.05, focusing on GO: BP. We report up to the top 10 significant biological processes with the largest odds ratios for select comparisons with n > 3 DEGs.

### Behavioral Testing

#### Open field test

At 3 days post injury (dpi), we placed individual mice in an open field chamber (12”x15”) containing no objects and allowed them to naturally explore for 15 minutes. We analyzed how much time the animal spent in corner, outer, inner, and center areas, as well as total ambulation using previously described automated methods [64]. We used n = 41-44 animals per experimental condition.

#### Novel object recognition

At 3 days post-injury (dpi), we placed the mice in the open field chamber for 15 minutes as described in *Methods: Open field test* to habituate them to the chamber. We then removed the mouse for 5 minutes and lightly cleaned the chamber using 70% ethanol. We then placed two identical objects in the chamber for training. Training consisted of two 10-minute trials in which the mouse was placed back in the chamber to familiarize themselves with the objects. The following day at 4 dpi, we replaced one of the familiar objects with a novel object of similar size. We placed the mouse back in the chamber for 5 minutes to freely explore the two objects. All cognitive assessment occurred during the light cycle under similar lighting and temperature conditions and was performed by the same experimenter. We tracked the mice’s exploration and the time they spent interacting with the two objects using previously described automated methods described in [64]. Since mice exhibit an innate preference for novelty, increased time spent with the novel object suggests that they recall which object is novel versus familiar. We expressed the novel object preference as the time spent interacting with the novel object divided by the total time spent interacting with either object. Any mouse with a sum total interaction time with either object <10 seconds was excluded from analysis [65] for a total n = 20-33 per experimental condition (Fig 1B), consistent with [4].

#### Contextual fear conditioning

At 8 days post-injury (dpi), we placed the mice in the fear conditioning chamber with sound attenuation for a total of 6 minutes, administering three 2-second, 0.4-mA shocks at 3 min, 4 min, and 5 min. One day later, at 9 dpi, we returned the mouse to the fear conditioning chamber for testing, recording their exploration for 3 minutes. We removed odor cues with 10% bleach and 70% ethanol between trials. An experimenter blinded to the experimental condition manually scored videos to determine the amount of time each animal spent freezing during the testing trial; the freeze percentage reflects the animal’s freezing duration over the total trial duration. Contextual fear conditioning was performed by the same experimenter at the same time of day (morning) during the light cycle. We used n = 35-42 animals per experimental condition, consistent with [4].

### Identification of miRNAs associated with novelty memory

To identify miRNAs associated with novelty memory, we first constructed a weighted average of the novel object preference (NOP) based on the volume of serum collected from each mouse in a particular pooled sample (see: Methods: GluA1/2+ extracellular vesicle isolation for pooled serum). This process allowed us to consolidate the behavior score in the same fashion as we consolidated serum EVs’ miRNAs while avoiding artificial inflation of the sample size. We then performed linear regression between the log2 read count of a single miRNA against the weighted novel object preference across serum samples, correcting for multiple comparisons using an FDR < 0.1. Next, we performed Least Absolute Shrinkage and Selection Operator (LASSO) regression on the entire dataset using *glmnet* for R with 10-fold cross-validation to identify an optimal panel of miRNAs that minimized the mean average error (MAE) while minimizing the risk of overfitting [66]. In miRNA panels derived from LASSO, conventional Pearson correlation p-values are not valid because they ignore the feature-selection process. Therefore, to validate our LASSO model, we performed permutation testing using MAE as our permutation statistic, since that was the metric used to optimize our LASSO model. We randomly permuted the NOP values 1000 times, and repeated the complete LASSO modeling procedure, including 10-fold cross-validation and model selection, for each permutation. We calculated an empirical p-value as the proportion of permuted models with a cross-validated MAE less than or equal to the observed model, using the recommended correction for randomly sampled permutations [67].

### Quantitative polymerase chain reaction (qPCR)

To confirm our sequencing results, we performed quantitative polymerase chain reaction (qPCR) on selected miRNAs identified as differentially expressed by sequencing analysis. We used the miRCURY LNA SYBR Green System (Qiagen) to prepare samples according to manufacturer instructions. qPCR was performed on the CFX384 Real-Time PCR System (Bio-Rad) and processed using the accompanying Maestro software (Bio-Rad). We ran each sample from sequencing in triplicate and used interplate calibrators (UniSp3, UniSp6) to ensure consistency across PCR plates. We confirmed sequencing findings in 21 unique miRNAs using qPCR, and found a Pearson’s correlation r = 0.55 (Fig S2).

### Cytokine analysis

We used the LEGENDplex Mouse Inflammation Panel (BioLegend) according to manufacturer’s instructions at the Penn Cytomics and Cell Sorting Shared Resource Laboratory to evaluate the presence of cytokines in our serum samples at 9 dpi. The panel included IL-23, IL-1α, IFN-γ, TNF-α, CCL2 (MCP-1), IL-12p70, IL-1β, IL-10, IL-6, IL-27, IL-17A, IFN-β, and GM-CSF. We ran n = 6-7 unpooled serum samples/experimental condition in duplicate for each cytokine and reported the average of the two concentrations. Each serum sample originated from a single animal. We excluded samples below BioLegend’s stated limit of detection and statistical outliers using the Robust Outlier detection Test (ROUT) (Q = 1%).

### Statistical analysis

Results are expressed as mean ± SEM. Statistical testing on miRNA sequencing data was performed using R (Version 4.4.1). Statistical testing for cognitive assessment was performed using GraphPad Prism Version 10 for Windows. Wald’s test was performed to identify differentially expressed miRNAs among experimental conditions, correcting for multiple comparisons using false discovery rate (FDR), with Q < 0.1 designating a discovery. Additional family-wise error rate correction (e.g., Sidak correction) was not applied, as FDR control is considered a standard and sufficient approach for genome-wide discovery analyses. Hypergeometric testing was performed to identify functionally enriched pathways for mRNAs targeted by miRNAs, controlling for multiple comparisons using FDR and Q < 0.1. We used logistic regression to classify experimental conditions and generate AUCs, and DeLong’s Method to generate confidence intervals for AUCs for EV miRNA biomarkers. For snRNA-seq data, we also designated discoveries with Q < 0.1. We used Fisher’s exact test to identify significantly enriched Gene Ontology (GO) Biological Processes, reporting the top 5 processes. ANOVA was used to compare behavioral performance and cytokine expression among experimental conditions, correcting for multiple comparisons using Sidak’s test and establishing significance at p < 0.05. ROUT was used to remove statistical outliers from cytokine analysis. Pearson correlation was performed to correlate between normally distributed log2 miRNA expression and behavior, as well as log2 miRNA expression from sequencing and qPCR.

## Results

### GluA1/2+ extracellular vesicles contain miRNA biomarkers indicative of injury and impact history

First, we investigated which miRNAs from GluA1/2+ EVs could serve as diagnostic biomarkers for concussive head injury. In our experimental model, we considered two concussion paradigms: concussion alone (unconditioned concussion) and concussion preconditioned by two subconcussive impacts delivered a minute apart (preconditioned concussion) (Fig 1A). Once cognitive assessment was complete at 9 days post-injury, we isolated serum from a terminal blood draw and used nanofluidic TENPO devices to isolate brain-enriched GluA1/2+ extracellular vesicles (EVs) for miRNA sequencing (Fig 1A), using serum pooled from 1-4 mice (mean = 2.1 mice/pooled serum sample) to produce the 500 μL required for consistent TENPO chip performance and efficient RNA extraction (Fig 1B).

At 9 days post-injury, miR-203-3p was the only miRNA that was significantly elevated in unconditioned concussion relative to sham (Fig 2A; Wald’s test p_adj_ = 0.07; log2FC = 1.7). When we increased the cumulative number of impacts by preconditioning the concussive impact with two subconcussive impacts, miR-203-3p remained significantly elevated (Fig 2A; Wald’s test p_adj_ = 0.007; log2FC = 2.2) and miR-205-5p also became significantly elevated (Fig 2A; Wald’s test p_adj_ = 0.08; log2FC = 1.9). Gene ontology (GO) enrichment analysis for biological processes (BP) for miR-203-3p and miR-205-5p revealed that the top terms involving these miRNAs included response to misfolded proteins, cytoskeletal and synaptic organization, and inflammation (Fig 2B; full list in Table S1).

**Fig 2.**
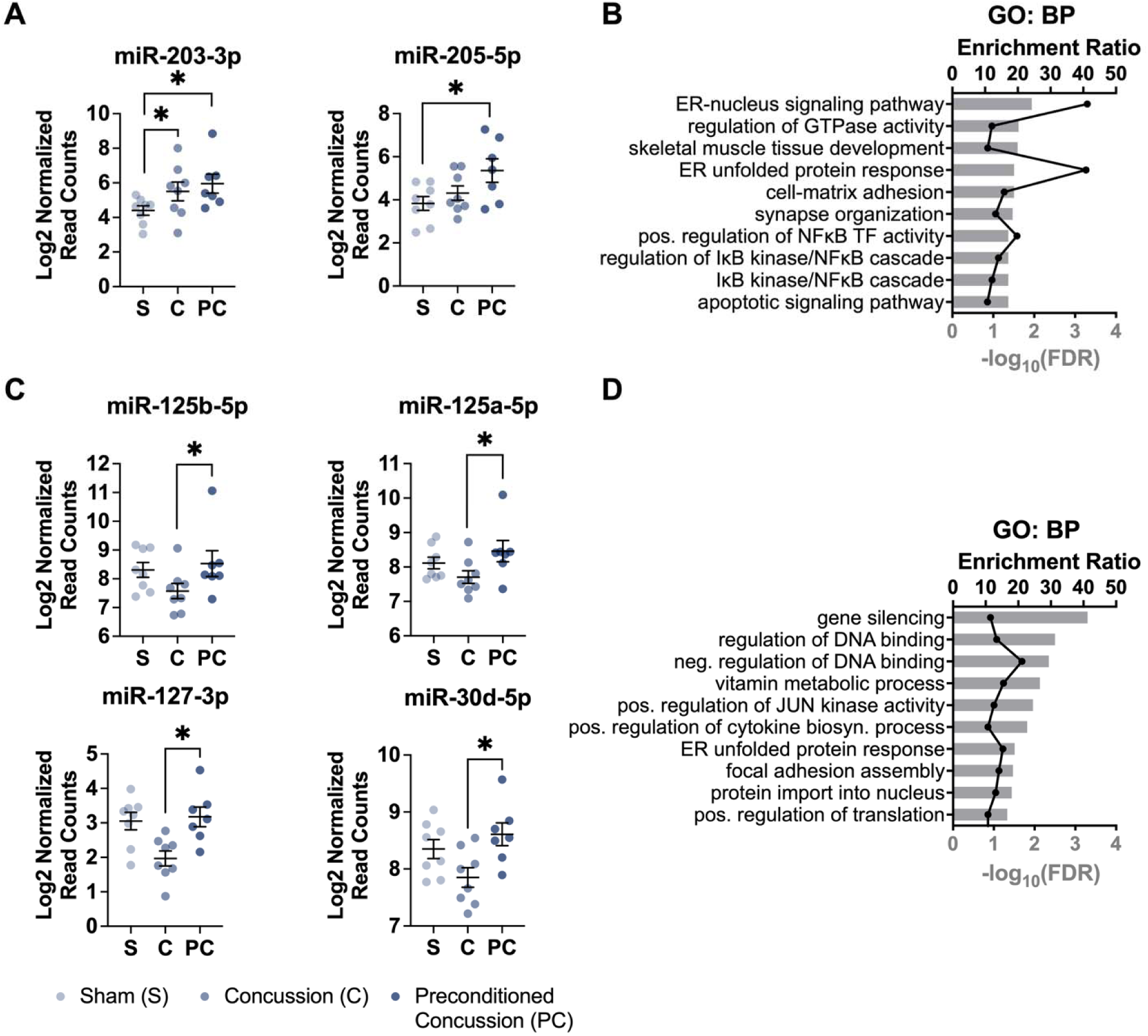
GluA1/2+ EV miRNA expression is dependent on head injury history. (A) GluA1/2+ EV miRNAs differentially expressed based on injury condition relative to sham for vehicle-treated mice – diagnostic miRNAs. (B) Top 10 Gene Ontology: Biological Processes (GO: BP) associated with diagnostic miRNAs. (C) GluA1/2+ EV miRNAs differentially expressed based between the two injury conditions for vehicle-treated mice – injury history miRNAs. (D) Top 10 GO: BP associated with injury history miRNAs. Data represented as mean +/- SEM. * FDR < 0.1. Each datapoint represents a pooled serum sample. Sham (S): N = 8, Concussion (C): N = 8, Preconditioned Concussion (PC): N = 7.

Next, we identified miRNAs from GluA1/2+ EVs which differed between unconditioned and preconditioned concussion. miR-125b-5p, miR-125a-5p, miR-127-3p, and miR-30d-5p were all significantly elevated for preconditioned concussion relative to concussion without prior impact history (Fig 2C; Wald’s test p_adj_ = 0.04, 0.09, 0.05, and 0.04; log2FC = 1.3, 0.9, 1.4, and 0.8, respectively). Notably, these miRNAs were distinct from those miRNAs broadly associated with injury (Fig 2A). GO: BP terms indicated that preconditioning induced changes in transcriptional and epigenetic regulation and stress/inflammatory signaling (Fig 2D; full list in Table S2).

We next assessed these differentially expressed miRNAs’ individual discriminative performance as candidate biomarkers by fitting separate logistic regression models for each miRNA to determine whether they could discriminate between binary experimental conditions. Within the discovery cohort, individual diagnostic miRNAs for either concussion or preconditioned concussion (Fig 2A), yielded AUCs between 0.73 and 0.86 for classification against sham. However, only the diagnostic miRNAs for preconditioned concussion versus sham, miR-203-3p and miR-205-5p, had confidence intervals that excluded AUC of 0.5 (95% CI: miR-203-3p, concussion vs sham: [0.45, 0.1], miR-203-3p, preconditioned concussion vs sham: [0.65, 1], miR-205-5p, preconditioned concussion vs sham: [0.57, 1] DeLong’s Method; Fig 3A), indicating discrimination above random guessing. Similarly, the miRNAs which independently discriminated preconditioned concussion from concussion (Fig 2C) had AUCs ranging from 0.79 to 0.93 (Fig 3A). All were superior to random guessing (95% CIs: miR-125b-5p: [0.57, 1], miR-125a-5p: [0.52, 1], miR-127-3p: [0.75, 1] and miR-30d-5p: [0.69, 1], DeLong’s Method), indicating their potential utility as impact history biomarkers, but they should be independently validated in future studies.

**Fig 3.**
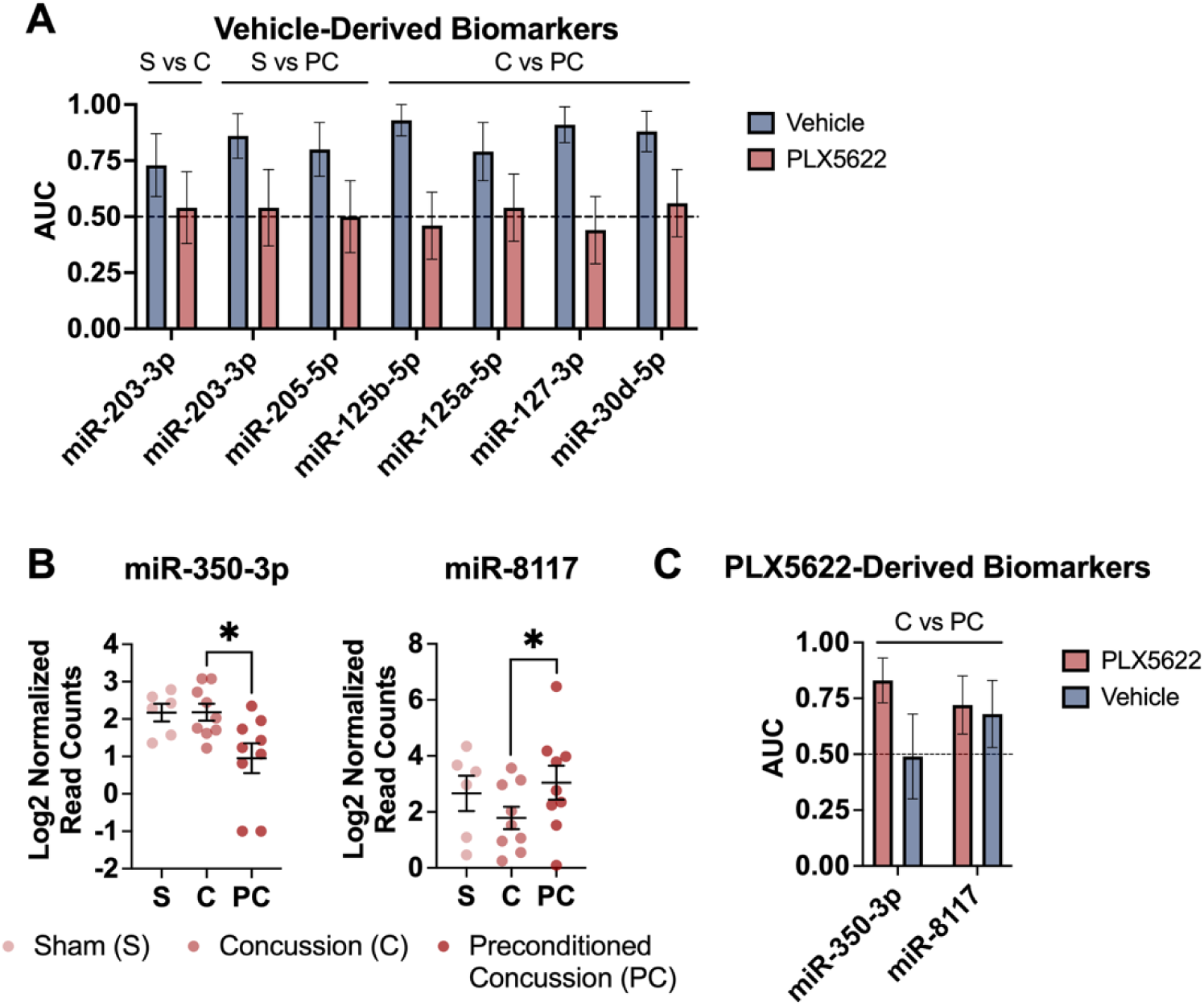
Microglial depletion modifies head injury history biomarkers. (A) Vehicle-derived GluA1/2+ EV miRNA biomarkers do not generalize to PLX5622-treated mice. (B) GluA1/2+ EV miRNAs differentially expressed based on injury condition for PLX5622-treated mice. * FDR < 0.1. (C) PLX5622-derived GluA1/2+ EV miRNA biomarkers miR-350-3p and miR-8117 do not generalize to vehicle-treated mice, but miR-8117 is comparable across treatments. Data represented as mean +/- SEM. Each datapoint represents a pooled serum sample. Sham (S), PLX5622: N = 6, Concussion (C), PLX5622: N = 9, Preconditioned Concussion (PC), PLX5622: N = 9.

### Microglia affect GluA1/2+ extracellular vesicle biomarkers and their diagnostic capacity

Past work shows that microglia play a key role in supporting and modifying neuronal function, suggesting that isolation of injury-related inflammatory miRNAs could help in the identification of differential recovery and repair phenotypes post-injury. Our GluA1/2+ EV signatures showed that preconditioned concussion led to a persisting expression of inflammation-mediated pathways compared to unconditioned concussion (Fig 2D). Relatedly, our prior work demonstrated that unconditioned concussion elicits an acute pro-inflammatory microglial response that preconditioning actively suppresses [4]. Collectively, this evidence led us to hypothesize that differential microglial activation acutely post-injury would affect EV biomarker signatures and efficacy.

To modulate the acute inflammatory response, we treated the mice with PLX5622, a brain-penetrant CSF1R-inhibitor that depletes microglia and other myeloid cells, to ensure near-complete depletion at the time of concussive injury (Fig S1). Immediately following injury, mice were returned to standard chow to permit microglial repopulation during the post-injury period. Consequently, the observed molecular and behavioral outcomes likely reflect the combined effects of microglial depletion at the time of injury and the subsequent repopulation process, rather than either phase in isolation. As before, we collected serum at 9 days post-injury and isolated GluA1/2+ EV miRNAs to test whether the diagnostic and impact history biomarkers we previously identified (Fig 2A) were altered following transient depletion of microglia and other CSF1R-dependent myeloid cells. To this end, we used the same logistic regression models trained on vehicle data and applied them to an independent test set of PLX5622 data. Following microglial depletion with PLX5622, neither diagnostic biomarker from the vehicle conditions (miR-203-3p, miR-205-5p; concussion versus sham or preconditioned concussion versus sham; Fig 2A) could discriminate sham from head injury, failing as diagnostic biomarkers for the PLX5622 conditions (AUC = 0.50 to 0.54; 95% CI: miR-203-3p, concussion vs sham: [0.21, 0.86]; miR-203-3p, preconditioned concussion vs sham: [0.20, 0.87]; miR-205-5p, preconditioned concussion vs sham: [0.18, 0.82], Fig 3A). Similarly, the injury history biomarkers from the vehicle conditions (miR-125b-5p, miR-125a-5p, miR-127-3p, and miR-30d-5p; concussion versus preconditioned concussion; Fig 2C) were no better than random guessing following microglial depletion (AUC = 0.44 to 0.56; 95% CIs: miR-125b-5p: [0.17, 0.74], miR-125a-5p: [0.26, 0.83], miR-127-3p: [0.16, 0.73] and miR-30d-5p: [0.26, 0.85], DeLong’s Method, Fig 3A). At 9 days post-injury, both diagnostic and injury history biomarkers lost their discriminative capacity following transient depletion and repletion of microglia and other CSF1R-dependent myeloid cells during acute injury.

We next considered the converse: whether there were GluA1/2+ EV miRNA biomarkers from PLX5622-treated animals (i.e., microglia-independent, injury-dependent) that could generalize to the vehicle condition. We identified no differentially expressed miRNAs between either injury group and sham following PLX5622 treatment. However, there were two miRNAs from the PLX5622-treated mice which were differentially expressed based on injury history: miR-350-3p and miR-8117 (Fig 3B; Wald’s test p_adj_ = 0.1 and 0.03; log2FC = -1.2 and 2.1, respectively). These differentially expressed miRNAs were also able to discriminate between injury histories in the PLX5622 condition with AUCs of 0.83 and 0.72, respectively (Fig 3C), although only miR-350-3p was superior to random guessing (95% CI: miR-350-3p: [0.63, 1], miR-8117: [0.46, 0.98], DeLong’s Method). However, when using the vehicle miRNA expression as an independent test set, neither miRNA’s discriminatory ability in the PLX5622 condition generalized to the vehicle treatment (miR-350-3p: AUC = 0.49, 95% CI [0.13, 0.86], miR-8117: AUC = 0.68, 95% CI [0.37, 0.98]; DeLong’s Method; Fig 3C). There were no significant GO: BP terms influenced by the combination of these two miRNAs.

We next investigated more directly how microglial depletion at the time of injury affected EV miRNA signatures, shifting our focus from identifying injury biomarkers to a more fundamental question of how transient depletion of microglia and other CSF1R-dependent myeloid cells affects neuron-enriched GluA1/2+ EV miRNA content within the same dataset. As a reference measure, there were no differentially expressed miRNAs in sham animals fed either vehicle or PLX5622 chow, indicating microglia depletion and repletion had no significant effects on neuron-enriched GluA1/2+ EV content. However, within concussed mice, 12 miRNAs were differentially expressed on the basis of microglia presence or absence: miR-127-3p, miR-30d-5p, miR-136-3p, miR-215-5p, miR-195a-5p, miR-381-3p, miR-199a-3p, miR-192-5p, miR-200a-3p, miR-122-5p, miR-379-5p, miR-194-5p (Fig 4A; Wald’s test p_adj_ ranged 0.0003 for miR-122-5p to 0.1 for miR-381-3p; log2FC ranged from 0.7 for miR-30d-5p to 2.63 for miR-122-5p). These miRNAs were associated with GO: BP terms relating to metabolic and homeostatic processes, proliferation, and apoptosis; a subset of the significant pathways are shown in Fig 4B; full list in Table S3. Notably, two of these miRNAs, miR-127-3p and miR-30d-5p, were shared among those affected by injury history (Fig 2C), raising the possibility of some shared mechanisms. There were no differentially expressed miRNAs in the preconditioned concussion condition for vehicle compared to PLX5622, as in sham.

**Fig 4.**
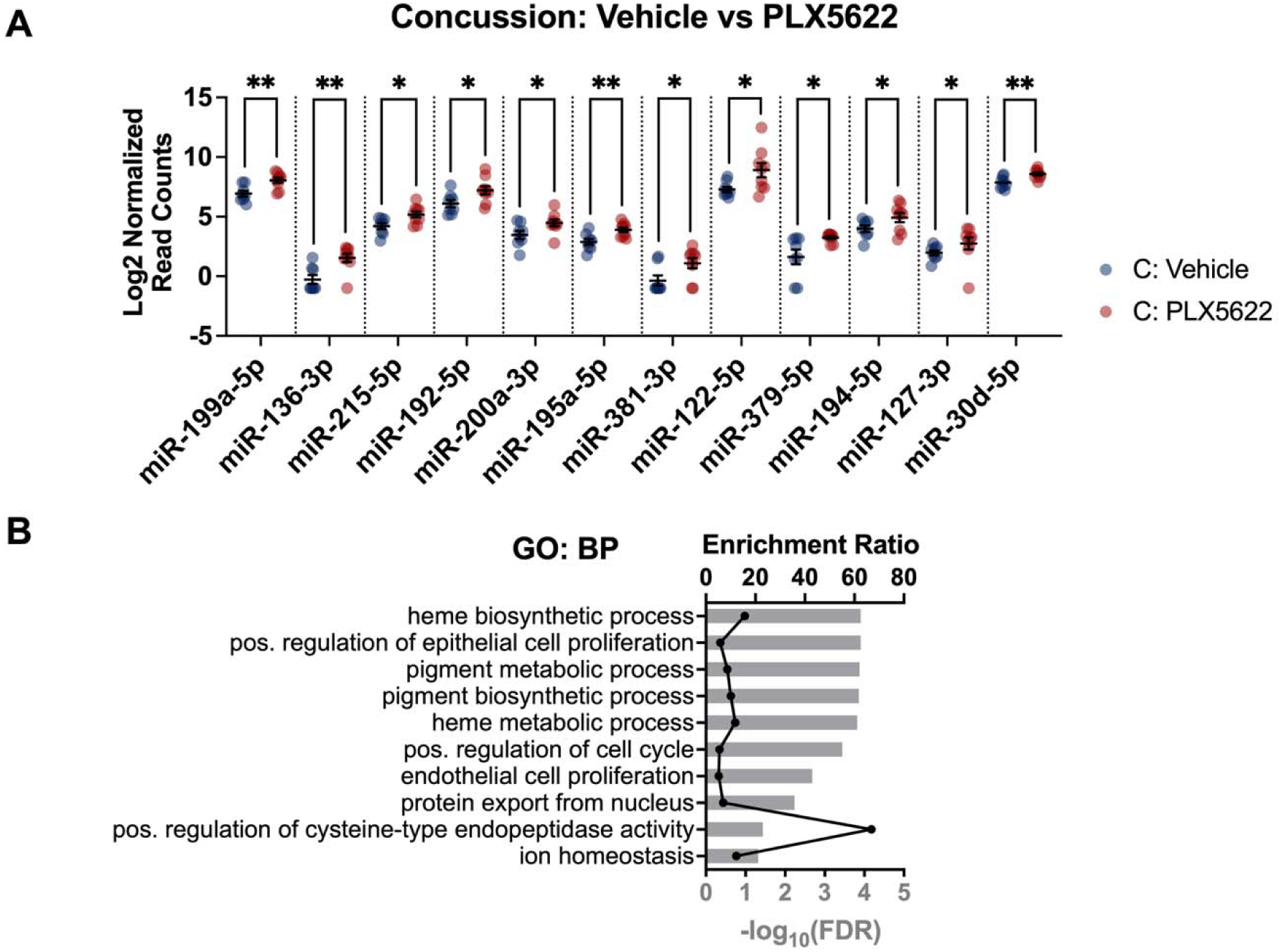
Microglial depletion changes GluA1/2+ EV miRNA expression relative to sham in concussed mice only. (A) GluA1/2+ EV miRNAs differentially expressed between vehicle and PLX5622 in concussed mice. * FDR < 0.1; ** FDR < 0.01. (B) Top 10 GO: BP associated with injury history miRNAs. Complete list can be found in Table S3. Data represented as mean +/- SEM. Each datapoint represents a pooled serum sample. Vehicle Concussion (C): N = 8, PLX5622 Concussion (C): N = 9.

Together, these findings demonstrate that GluA1/2+ EV miRNA signatures at 9 days post-injury are shaped by transient depletion and repletion of microglia and other CSF1R-dependent myeloid cells during the acute injury, and warrant future investigation in an independent test set.

### Single-nuclei transcriptomics reveals cellular substrates of differential injury and microglial depletion

Neuron-enriched GluA1/2+ EV signatures revealed divergent molecular trajectories related to injury and impact history and microglial dependence. To characterize the neuronal transcriptional consequences of microglial depletion and repopulation, we used an independent molecular approach: single-nuclei RNA-seq (snRNA-seq) of brain tissue from the injured hemisphere at 9 days post-injury. We selected snRNA-seq because of its suitability for isolating neuronal RNA, the primary originating cells of GluA1/2+ EVs. Neurons can be difficult to dissociate intact for single-cell RNA-seq (scRNA-seq) [68], and are thus better suited to snRNA-seq; however, by choosing snRNA-seq, we sacrificed a direct readout of repopulating microglia, which are better isolated and characterized with scRNA-seq [69]. Instead, microglial contributions were inferred from the effects of PLX5622-mediated depletion and repopulation on neuronal gene expression, complementing our previous bulk RNA sequencing analysis of isolated microglia from the same experimental model [4].

Among our nuclei, we identified 10 neuronal clusters and 1 cluster of oligodendrocytes/oligodendrocyte precursor cells (OPCs) (Fig 5A; Fig S3A), with similar nuclei type distributions and counts across experimental conditions (Fig S3B). To identify nuclei types in our dataset, we harmonized leiden clusters with annotations derived from the Allen Mouse Brain Cell Atlas, leading us to exclude a cluster of 3162 nuclei (of the 21476 nuclei that originally passed quality control) for which we could not confidently ascribe a cellular origin.

**Fig 5.**
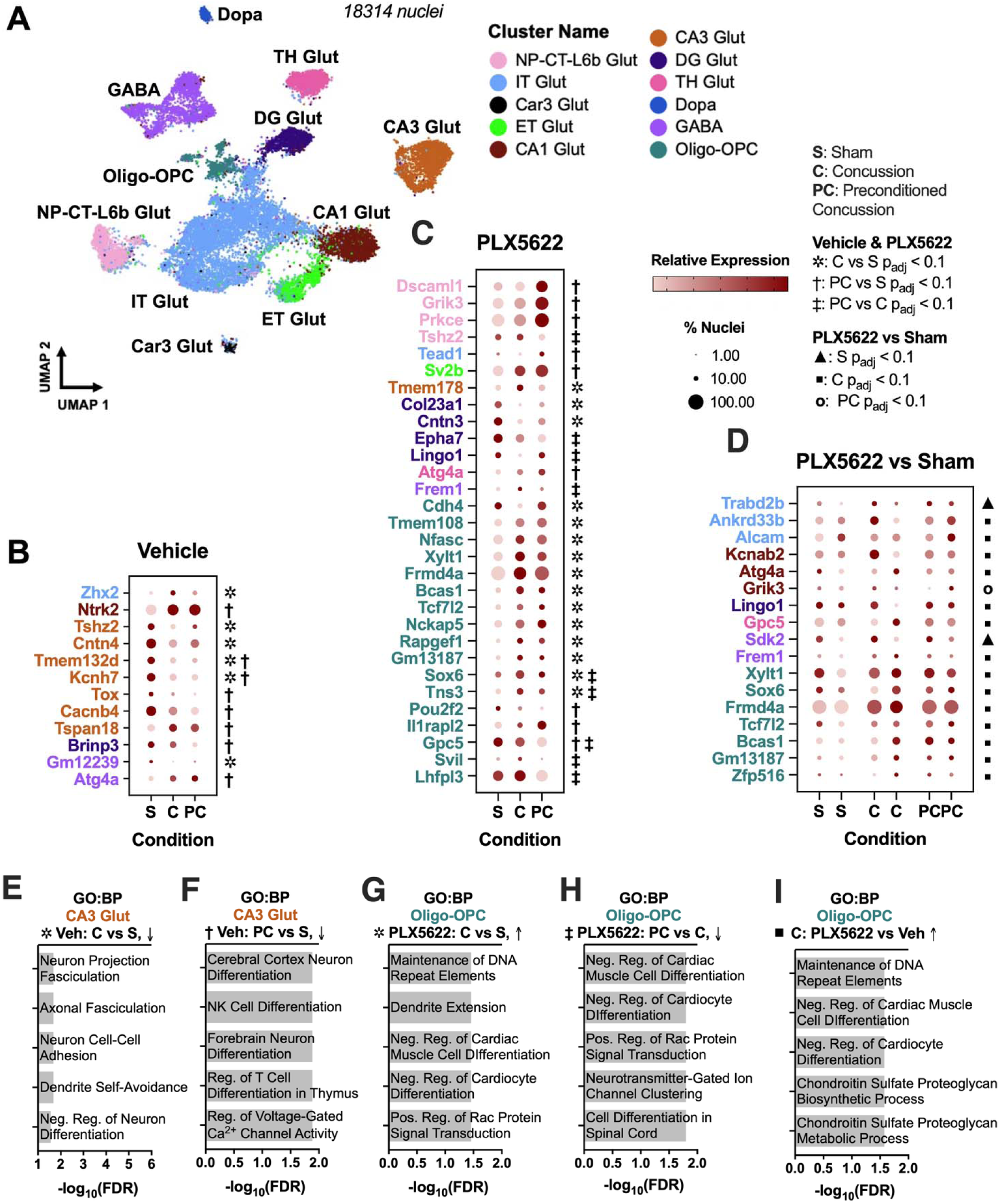
snRNA-seq reveals injury- and inflammation-dependent transcriptomic changes to CA3 and oligodendrocytes/OPCs. (A) UMAP of nuclei pooled from all experimental conditions, showing 11 clusters, including 10 neuronal subtypes and 1 oligodendrocyte-OPC cluster. (B-D) Dot plots showing differentially expressed genes (DEGs) across injury conditions (B) within the vehicle treatment group, (C) within the PLX5622 treatment group, and (D) comparing vehicle and PLX5622 treatment groups within each injury condition. Gene font color denotes the nuclei type, dot color indicates relative expression level, and dot size indicates the percentage of nuclei in the cluster that express that gene. (E-I) Up to 10 significant Gene Ontology: Biological Processes (GO: BP) terms for select sets of DEGs (↑ = upregulated, ↓ = downregulated): (E) CA3 Glut genes downregulated by vehicle-concussion relative to vehicle-sham, (F) CA3 Glut genes downregulated by vehicle-preconditioned concussion relative to vehicle-sham, (G) Oligo-OPC genes upregulated by PLX5622-concussion relative to PLX5622-sham, (H) Oligo-OPC genes downregulated by PLX5622-preconditioned concussion relative to PLX5622-concussion, and (I) Oligo-OPC genes upregulated by PLX5622-concussion relative to vehicle-concussion. NP-CT-L6b Glut: Near-projecting-corticothalamic-Layer 6b Glutamatergic Neurons; IT Glut: Intratelencephalic Glutamatergic Neurons; Car3 Glut: Car3+ Glutamatergic Neurons; ET Glut: Extratelencephalic Glutamatergic Neurons; CA1 Glut: CA1 Hippocampal Glutamatergic Neurons; CA3 Hippocampal Glutamatergic Neurons; DG Glut: Dentate Gyrus Glutamatergic Neurons; TH Glut: Thalamic Glutamatergic Neurons; Dopa: Dopaminergic Neurons; GABA: GABAergic Neurons; Oligo-OPC: Oligodendrocyte-Oligodendrocyte Precursor Cells. For Vehicle and PLX5622 Treatment: ✲: C vs S p_adj_ < 0.1; †: PC vs S p_adj_ < 0.1; ‡: PC vs C p_adj_ < 0.1; For Vehicle vs. PLX5622 Treatment: ▴: S p_adj_ < 0.1; ▪□: C p_adj_ < 0.1; ο: PC p_adj_ < 0.1. S: Sham; C: Concussion; PC: Preconditioned Concussion. N = 4 animals/condition; nNuclei = 2637 (Vehicle-Sham), 3492 (Vehicle-Concussion), 3022 (Vehicle-Preconditioned Concussion), 3100 (PLX5622-Sham), 2947 (PLX5622-Concussion), 3116 (PLX5622-Preconditioned Concussion).

These omitted nuclei likely consisted of a mix of astrocytes, other glia, and rare neuron subtypes. Among the 18314 successfully annotated nuclei, examined differentially expressed genes (DEGs) (Fig 5B-D) and the corresponding pathways using over-representation analysis based on up- or down-regulation of individual DEGs (Fig 5E-I). First, we examined DEGs across vehicle-treated injury conditions (Fig 5B, all DEGs listed; Fig S4 for volcano plots; Table S4 for p_adj_ and log2FC). For both unconditioned and preconditioned concussion, CA3 glutamatergic neurons exhibited the most changes in gene expression relative to sham, sharing two genes (*Tmem132d* and *Kcnh7*) in common. In CA3 glutamatergic neurons, concussion resulted in downregulation of genes related to neurite organization and neuronal adhesion (Fig 5E), and preconditioned concussion downregulated genes relating to neuron differentiation and ion channel activity (Fig 5F). Among the injury-associated DEGs, only *Cacnb4* is a target of our diagnostic GluA1/2+ EV miRNAs — specifically, miR-205-5p for preconditioned concussion (Fig 2A). There were no DEGs for unconditioned versus preconditioned concussion. Among our measured nuclei types, CA3 glutamatergic neurons experienced the most transcriptional changes following concussive head injury, and these changes were similar regardless of prior impact history and were not targeted by injury-associated GluA1/2+ EV miRNAs.

We next considered how transient depletion and repletion of microglia and other CSF1R-dependent myeloid cells with PLX5622 affected the injury response. Oligodendrocytes/OPCs, which possess immune-related and microglia-dependent functions [70], were prominently affected by injury after microglial depletion (Fig 5C, all DEGs listed; Fig S5 for volcano plots; Table S4 for p_adj_ and log2FC). Two genes (*Sox6, Tns3*) were upregulated by concussion relative to both sham and preconditioned concussion; *Sox6* was implicated in oligodendrocyte differentiation and *Tns3* was implicated in Rac Protein Signal Transduction, controlling actin cytoskeleton reorganization for myelin ensheathment. Only *Gpc5*, a heparan sulfate proteoglycan produced by OPCs, was injury-history independent. These functions relating to differentiation, structural organization, and extracellular matrix are reflected in the GO: BP Analysis for genes upregulated by concussion following PLX5622 treatment (Fig 5G-H). None of these genes, regardless of nucleus type, were regulated by differentially expressed GluA1/2+ EV miRNAs (Fig 3B). Together, these findings indicate that microglial depletion is associated with transcriptional programs relating to oligodendrocyte maturation and myelin sheath formation, but only for unconditioned concussion.

Finally, we considered how PLX5622 treatment affected gene response within an injury condition. Oligodendrocyte/OPCs were the most affected nucleus type and concussion was the most affected injury condition (Fig 5D, all DEGs listed; Fig S6 for volcano plots; Table S4 for p_adj_ and log2FC). Again, however, none of these genes were among known targets of GluA1/2+ EV miRNAs (Fig 4A). The genes upregulated by PLX5622 treatment in the concussion condition compared to vehicle treatment, such as *Sox6* as above, were involved in oligodendrocyte differentiation (*Sox6)* and chondroitin sulfate proteoglycan synthesis (*Xylt1*) (Fig 5I). These functions support oligodendrocytes’ ability to myelinate and remodel the extracellular environment to support myelination.

Together, the snRNA-seq data identify CA3 glutamatergic neurons’ transcriptional programs as persistently affected by concussion regardless of injury history, and reveal that microglial depletion is associated with transcriptional signatures suggestive of oligodendrocyte maturation and remyelination in unconditioned concussion — a cellular mechanism consistent with the unimpaired behavior observed in these animals compared to preconditioned concussion.

### GluA1/2+ extracellular vesicle miRNA signatures correlate with acute cognitive recovery

We recently showed that subconcussive preconditioning, induced by delivering two subconcussive impacts immediately before a concussive impact, eliminates concussion-associated cognitive deficits in novel object recognition [4]. Here, we tested whether this protective effect, and the cognitive deficit it prevents, persisted in these experimental conditions and whether any behavioral change is microglia-dependent. Using an independent cohort of mice maintained on the semi-synthetic diet required for PLX5622 administration [71], we first aimed to confirm the original behavioral phenotype. However, in vehicle-treated mice, unconditioned concussion caused a non-significant decrease in novel object recognition relative to sham (Fig 6A; two-way ANOVA with Sidak correction: p = 0.11, fold-change (FC) = 0.9, Cohen’s *d* = 0.47), whereas preconditioned concussion in vehicle-fed mice had no novel object recognition deficit (Fig 6A), only partially reproducing our earlier finding [4].

**Fig 6.**
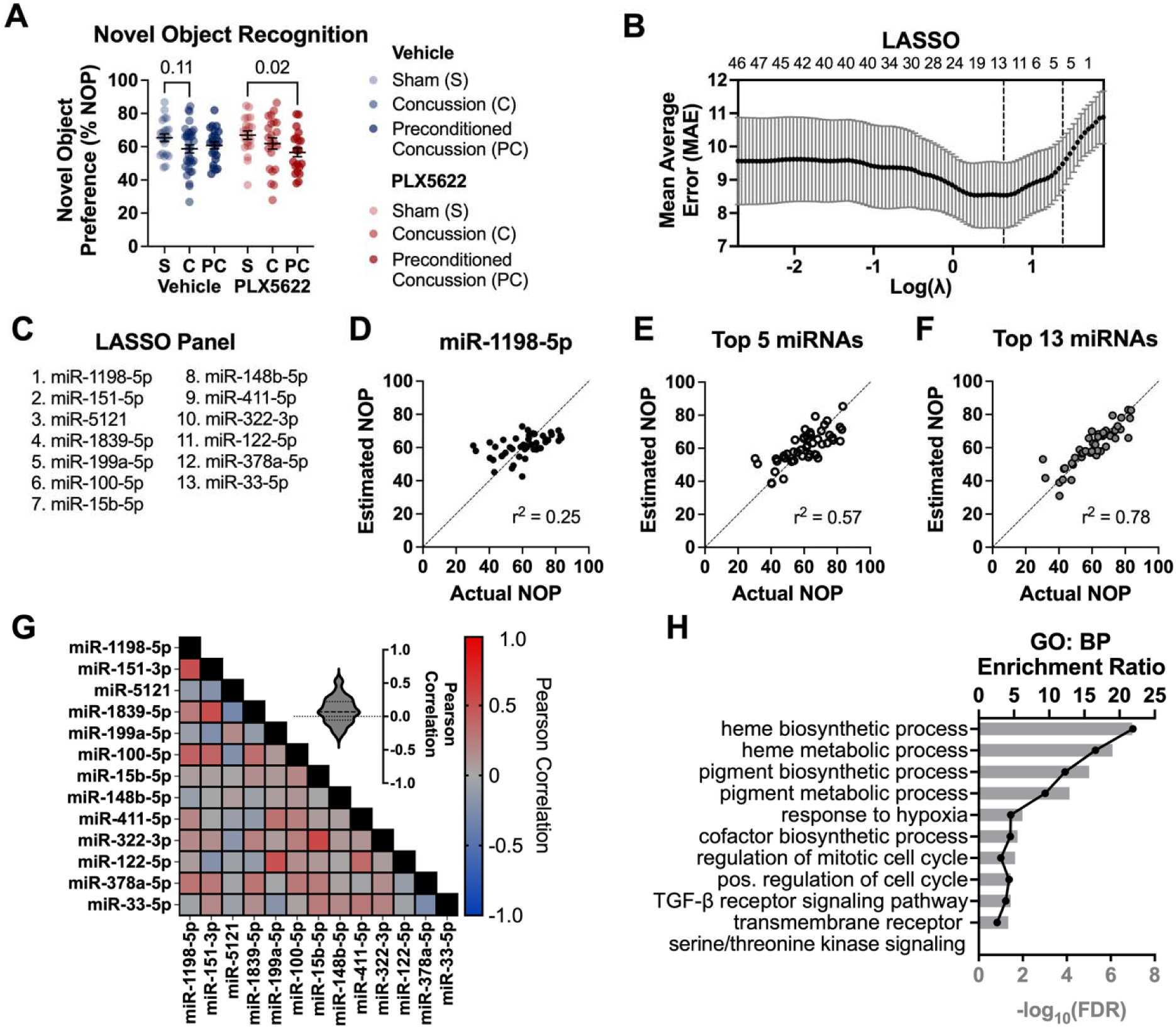
GluA1/2+ EV miRNAs reflect injury- and inflammation-dependent changes in novel object recognition. (A) PLX5622 treatment prior to injury introduces a significant deficit (p < 0.05) in novel object recognition (NOR) for preconditioned concussion that is not present with vehicle treatment. Two-way ANOVA with Sidak’s test for multiple comparisons. Each datapoint represents a single animal. (N = 20-33/condition - see: Fig 1) (B) Using 13 miRNA predictors minimizes the error of novel object preference estimation. (C) Ordered list of LASSO panel (D-F) Increasing the number of novel object preference (NOP) predictors from (D) the top predictor generated by LASSO (miR-1198-5p) to (E) the top 5 and (F) the top 13 increases r^2^ from 0.25 to 0.57 to 0.78 using Pearson’s correlation. Each datapoint represents a pooled serum sample and its aggregate NOP score (N = 47 total). (G) Each of the top 13 predictor miRNAs are uncorrelated with the others. Pearson’s correlation. (H) Top 10 GO: BP terms for the top 13 miRNAs. Complete list with FDR < 0.1 can be found in Table S5. Data represented as mean +/- SEM. NOP: Novel object preference. S: Sham; C: Concussion; PC: Preconditioned Concussion.

We next considered how transient depletion of microglia and other CSF1R-dependent myeloid cells affected novel object recognition. Importantly, in sham animals, microglial depletion had no effect on novel object recognition (Fig 6A). In comparison, concussed mice with microglial depletion demonstrated no significant deficit in novel object recognition. Strikingly, preconditioned concussion, which exhibited no novel object recognition deficits in the vehicle condition, had a significant novel object recognition deficit following microglial depletion (Fig 6A; two-way ANOVA with Sidak correction: p = 0.02, FC = 0.85, Cohen’s *d* = 0.87).

Having shown that GluA1/2+ EV miRNAs reflect microglia-dependent injury biology, we last asked whether EV miRNA content correlates with acute cognitive outcomes. By collecting GluA1/2+ EVs at 9 days post-injury, we were able to directly correlate novel object recognition performance with GluA1/2+ EV content in the same mice (Fig 1, 6). Notably, serum samples were pooled according to similarity in individual novel object preference (NOP) scores, and a weighted average of the individual NOPs was used to represent each pooled sample in the correlation analysis (see Methods: Identification of miRNAs associated with novelty memory), thereby avoiding pseudoreplication.

First, we considered whether there were miRNAs that correlated with prior cognitive outcome regardless of injury or microglia depletion. miR-1198-5p and miR-151-3p each independently showed a modest correlation with aggregate novel object recognition (miR-1198-5p: r^2^ = 0.25, p_adj_ = 0.07; miR-151-3p: r^2^ = 0.24, p_adj_ = 0.07, Pearson’s correlation with FDR correction), but no individual miRNAs significantly correlated with contextual fear conditioning. We next considered whether using a panel of miRNAs could improve correlation with novel object preference. We performed LASSO using 10-fold cross-validation to identify such a panel (Fig 6B), listed in order in Fig 6C. We focused on three different predictor models: (1) a single miRNA (miR-1198-5p), selected to minimize the prediction error (empirical p = 0.43), (2) a panel of 5 miRNAs which balanced the model error while minimizing the risk of overfitting (empirical p = 0.042), and (3) a panel of 13 miRNAs which minimized the mean absolute error of the predictive model (empirical p = 0.017; Fig 6D-F; listed in order in Fig 6C). Unsurprisingly, miR-1198-5p and miR-151-3p were the top two miRNAs in the panel. We verified that the miRNAs in the panels were independent from each other: none of them had a correlation which exceeded 0.52 (Fig 6G). The correlation between the actual novel object preference and the estimated novel object preference using a 1-miRNA, 5-miRNA or 13-miRNA panel improved from r^2^ = 0.25 to 0.57 to 0.78 (Fig 6D-F). Although the 5-miRNA panel was not significantly associated with any GO: BP terms after FDR correction, the 13-miRNA panel was associated with GO: BP terms involved in oxidative stress and cell proliferation (abbreviated list in Fig 6H; complete list in Table S5). Due to our small sample size, we were not able to validate these findings in an independent test set, but mitigated overfitting by using cross-validation and performed permutation testing to verify model performance over random chance [72]. Together, these results suggest a possible rationale to identify miRNAs associated with cognitive outcome in future studies.

## Discussion

We set out to test microglia’s role in cognitive outcomes following concussion versus preconditioned concussion. The results were not what a simple, unidirectional model of neuroinflammation would predict. Transiently depleting microglia and other CSF1R-dependent myeloid cells before a single concussive impact had no measurable effect on cognitive performance, but depleting microglia before preconditioned concussion produced a cognitive deficit where none existed before, suggesting that the absence of a cognitive deficit normally seen after prior subconcussive exposure depends on microglia being present at the time of injury. Rather than acting as a uniform driver of harm or repair, microglia appear to play a role that is contingent on the brain’s injury history. These findings extend our earlier observation by showing that although preconditioning broadly dampens acute microglial reactivity [4], microglia are still required for an unimpaired outcome.

Some, but not all, of the effects of removing microglia in our study parallel findings in ischemic stroke. In ischemic preconditioning, microglia establish neuroprotection against subsequent ischemic events [3], and in unconditioned ischemia, acute microglial depletion often worsens stroke recovery [7]. More broadly, it is often the case in neurological disease that microglia are acutely protective before becoming chronically overactive and impeding recovery [73]. Our findings add an important dimension to this framework: our preconditioning model attenuates pro-inflammatory responses [4], yet transient depletion of microglia and other CSF1R-dependent myeloid cells worsens cognitive outcomes. This observation suggests that anti-inflammatory treatments in preclinical models may not be universally effective when modeling the effect of subconcussive insults prior to injury, an exposure profile common in contact sports.

Our data show that GluA1/2+ EV miRNA biomarkers, measured at 9 days post-injury, are influenced by transient depletion and repletion of microglia and other CSF1R-dependent myeloid cells. Both diagnostic miRNAs and injury history miRNAs lost their discriminatory capacity following microglial depletion, and 12 miRNAs were differentially expressed in concussed mice on the basis of microglial presence or absence. That these neuron-enriched EV biomarkers depend on microglia is itself informative: it suggests that GluA1/2+ EVs are capturing the neuronal response to the inflammatory milieu, not merely intrinsic neuronal injury.

This work builds on substantial efforts to develop blood biomarkers for TBI [11]. Protein biomarkers such as GFAP and UCH-L1 provide clear indication of acute injury and, to some extent, injury severity, while NfL and tau indicate chronic degeneration, with recent literature demonstrating NfL also has prognostic value [11–14]. These biomarkers measure common endpoint processes – neuronal death, axonal degeneration, glial reactivity – and are valuable for that reason. However, they do not identify the specific cellular mechanisms driving recovery and therefore have limited utility for guiding therapeutic intervention [15], particularly for mild injury, where cellular degeneration is less pronounced. In our model, we measured 13 common cytokines at 9 days post-injury and found no significant changes (Fig S7), consistent with the limited diagnostic value of circulating cytokines in the post-acute phase [17,74,75]. By contrast, functional enrichment analysis of GluA1/2+ EV miRNAs collected at the same timepoint indicate possibly ongoing inflammatory processes, suggesting that EV miRNAs provide a potentially more sensitive or persistent indicator of brain inflammation – or, alternatively, a longer-lasting molecular memory of the inflammatory challenge. This possibility warrants direct investigation. Some of the specific biomarkers we identified, such as miR-203-3p, align with those previously found in human TBI EV diagnostic panels [40], and inhibiting miR-203-3p improves TBI outcomes [76], supporting their biological relevance.

To complement our behavioral and EV biomarker findings, we used snRNA-seq as an independent molecular readout with cell-type specificity. Among vehicle-treated animals, CA3 glutamatergic neurons exhibited the most persistent changes in gene expression following concussion, regardless of preconditioning status. Changes in neurite organization, neuronal adhesion, and ion channel activity are consistent with CA3’s well-established vulnerability to TBI [77] and the synaptic plasticity and circuit-level dysfunction documented in prior work [78,79]. That CA3 changes were similar across injury histories suggests that preconditioning does not prevent the initial injury to neural circuits, but rather, engages microglia and potentially other myeloid cells [4]. The similarity in CA3 changes across injury histories also aligns with our behavioral findings in this study.

Transient depletion of microglia and other CSF1R-dependent myeloid cells revealed a distinct cellular transcriptional response in oligodendrocytes/OPCs. In unconditioned concussion, PLX5622 treatment was associated with upregulation of genes in oligodendrocytes/OPCs relating to differentiation (*Sox6*), myelin ensheathment (*Tns3*), and extracellular matrix remodeling (*Xylt1*). This pro-myelination signature provides a candidate mechanism for the absence of a deficit we observed in PLX5622-treated concussed mice: by removing the acute inflammatory response, transient depletion and repletion of microglia and other CSF1R-dependent myeloid cells may have permitted oligodendrocyte maturation and circuit repair that would otherwise be suppressed. This interpretation is consistent with work in ischemic stroke showing that transient acute microglial depletion supports oligodendrogenesis and remyelination [58]. In contrast, preconditioned concussion showed no significant changes with PLX5622 treatment in either the snRNA-seq or EV miRNA data, suggesting that the preconditioning response may engage different recovery mechanisms, including non-nuclear, post-transcriptional changes in unmeasured cell types, that do not rely on the same transcriptional programs for oligodendrocyte-mediated repair.

Although we attempted to connect the EV miRNA biomarkers directly to snRNA-seq gene expression changes, only *Cacnb4*, a voltage-gated calcium channel subunit involved in dendritic spine regulation [80], was shared between the two datasets. This limited overlap is expected: snRNA-seq measures nuclear RNA, which is less susceptible to miRNA regulation than cytoplasmic RNA [81], and the two approaches capture different molecular layers – EV miRNAs reflect intercellular signaling while snRNA-seq reflects intranuclear transcription. As such, the two approaches characterize complementary aspects of the injury response, converging on overlapping biological themes: synaptic reorganization, inflammatory cascades, and myelination. Both pro-inflammatory microglia and inflammation-mediated oxidative stress contribute to OPC maturation and myelin repair [50,82], linking the EV miRNA pathways to the snRNA-seq findings thematically even in the absence of direct gene-target overlap.

A final contribution of this work is the identification of an EV miRNA signature correlated to cognitive recovery. Using LASSO regression with 10-fold cross-validation, we identified a panel of up to 13 miRNAs that estimated prior novel object recognition performance with moderate-to-strong correlation (r^2^ = 0.78 for the full panel). Among the top miRNAs in the panel, the first four, including miR-1198-5p and miR-151-3p, were not significantly associated with a particular injury condition or inflammatory state and may represent miRNAs intrinsically associated with memory and cognition. Indeed, miR-1198-3p is predicted to regulate *Serca2* expression in the hippocampus, controlling neuronal calcium homeostasis and long-term potentiation [83], and miR-151-3p is upregulated by LTP induction in the dentate gyrus [84] and microglia-enriched miR-151-3p contributes to neuroprotection following spinal cord injury [85]. The entry of injury-and inflammation-dependent miRNAs (miR-199a-5p, miR-122-5p) later in the panel suggests a potentially hierarchical model: baseline cognitive capacity may be the primary driver of outcome, with injury and inflammation fine-tuning the recovery trajectory. This interpretation suggests that EV signatures could capture both the fundamental neurobiology of memory, as well as inflammatory modulation. Due to sample size limitations, we were unable to validate these findings in an independent test set or in female mice, and these results should be considered hypothesis-generating. Nonetheless, they establish a rationale for developing EV miRNA panels as prognostic biomarkers for cognitive recovery that capture the ongoing cellular processes that influence long-term outcome.

Several methodological considerations bear on the interpretation of our findings. First, we depleted microglia and other CSF1R-dependent myeloid cells with PLX5622 before injury and allowed them to repopulate during the recovery period, rather than maintaining depletion throughout. This design reflects our primary question: whether microglia present at the time of injury establish the cognitive trajectory. However, because microglial repopulation occurred during the post-injury period, our findings cannot distinguish the effects of microglial depletion at the time of injury from those of the subsequent repopulation process or the interaction between repopulating microglia and injury-induced responses. Rather, the observed behavioral and molecular outcomes likely reflect the combined consequences of transient depletion followed by post-injury repopulation. Repopulating microglia appear within 3 days of PLX5622 removal and largely normalize by 21 days [51,52], sharing similar surveillance activity and inflammatory gene expression with naïve microglia [86,87]. Given this timeline, our NOR assessment at 4 days post-injury primarily reflects the acute absence of microglia, while by 9 days post-injury (EV collection, snRNA-seq, fear conditioning), microglia are substantially repopulated. Our neurobehavioral findings are consistent with this interpretation: at 3-4 days post-injury, before significant repopulation, we observed decreased ambulation (Fig S8A), and no deficits in novel object recognition following TBI, consistent with depletion-phase effects [48,49]. It is possible that this decreased ambulation with PLX5622 treatment affected novel object recognition, however, because the decreased ambulation occurred independently of injury, we do not believe that it significantly influenced the novel object deficit in preconditioned mice following PLX5622 treatment, especially given that we observed no differences in NOR in sham animals between vehicle and PLX5622 treatment. Similarly, the injury-independent fear conditioning deficit at 9 days (Fig S8C) is consistent with repopulation-phase effects reported by others [59]. Continuous microglial depletion spanning the injury and recovery periods would be a valuable complementary experiment to disambiguate acute from chronic microglial contributions, but was beyond the scope of this study.

Second, CSF1R inhibition with PLX5622 affects peripheral monocytes and macrophages [88], which could influence the inflammatory cross-talk reflected in GluA1/2+ EV miRNAs at 9 days post-injury. Future studies using *Csf1r*^ΔFIRE/ΔFIRE^ mice in lieu of PLX5622 will be better equipped to isolate microglia-specific mechanisms. However, macrophage infiltration is delayed relative to the acute microglial response [89], making peripheral immune cells unlikely contributors to the preconditioning effect, which depends on impacts occurring within minutes [4]. Additionally, PLX5622 treatment combined with injury could affect the cellular origin of GluA1/2+ EVs, given notable *Gria2* expression in OPCs (Fig S3C) [49] and the profound PLX5622-dependent changes in oligodendrocyte/OPC gene expression we observed. Third, this study used only male mice, which exhibit a swifter and more pronounced neuroinflammatory profile acutely post-TBI [44]; our findings may not extend to female mice and these biomarker profiles, and the neuroinflammatory mechanisms they reflect, should be considered preliminary. Fourth, all animals were maintained on a semi-synthetic diet (AIN-76A) required for PLX5622 incorporation. We have previously noted that this diet does not affect the behavioral outcomes measured here, but does influence EV miRNA biomarker profiles [71], and the biomarker findings in this study should be interpreted within the context of this dietary background. Fifth, our biomarker panel was defined using serum collected after acute behavioral assessment, prohibiting true outcome prognosis, and using sample pooling required to meet volume standards. This experimental design was required due to the terminal nature of our blood draws in this preclinical model, but allowed us to identify hypothesis-generating, group-level biomarker associations that can now guide the selection of candidate analytes for future longitudinal, individually-resolved validation studies. Finally, these results should not be interpreted to mean that subconcussive exposure in humans could be considered protective, given the artificial preclinical scenario over short recovery windows explored here. Rather, the more clinically meaningful interpretation is that prior head impact history and inflammatory context may modify concussion outcome and associated biomarkers.

## Conclusions

This work reframes concussion recovery as a process influenced by microglia and shaped by prior exposure history. Microglial depletion had no effect on outcome after concussion alone but produced a cognitive deficit specifically after preconditioned concussion, suggesting that microglia are required for an unimpaired outcome only in the context of prior subconcussive exposure. This history-dependence challenges the assumption that post-concussive neuroinflammation is a single process with a single therapeutic target. Brain-derived GluA1/2+ EVs provide a blood-accessible readout of this biology, complementing protein biomarkers of cellular damage by instead reflecting how the brain is responding to injury. That a preliminary EV miRNA panel could predict cognitive outcome suggests these tools may eventually support more individualized approaches to concussion management — identifying not just who was injured, but who is likely to recover.

## List of abbreviations

EVs: Extracellular Vesicles
TBI: Traumatic Brain Injury
IACUC: Institutional Animal Care and Use Committee
dpi: days post-injury
IHC: Immunohistochemistry
TENPO: Track Etched NanoPOre
FDR: False Discovery Rate
GO: Gene Ontology
BP: Biological Processes
LASSO: Least Absolute Shrinkage and Selection Operator
qPCR: quantitative Polymerase Chain Reaction
snRNA-seq: single-nuclei RNA sequencing
scRNA-seq: single-cell RNA sequencing
ROUT: Robust Outlier detection Test
FC: Fold Change
OPCs: Oligodendrocyte Precursor Cells
DEGs: Differentially Expressed Genes

## Declarations

### Availability of data and materials

The sequencing datasets supporting the conclusions of this article are available in the NCBI Gene Expression Omnibus (GEO), Accession Number GSE330926 [https://www.ncbi.nlm.nih.gov/geo/query/acc.cgi?acc=GSE330926]. All other datasets are included within the article (and its additional files).

### Authors’ contributions

Conceptualization: EDA, DFM

Methodology: EDA, DVA, APG, TK, DAI, DFM

Investigation: EDA, DVA, TK

Formal Analysis: EDA, TK Data

Curation: EDA, DVA

Validation: EDA

Software: EDA

Visualization: EDA

Project Administration: APG

Supervision: DAI, DFM

Funding Acquisition: DAI, DFM

Writing—original draft: EDA, DFM

Writing—review & editing: EDA, DAI, DFM

## Acknowledgements

We thank Jean G. Rosario for his sequencing expertise.

## Additional Files

### SupplementalFigures.pdf

**Fig. S1. Validation of microglial depletion.** Representative coronal sections at -1.5-2 mm from bregma following 10d of (A) Vehicle or (B) PLX5622 treatment showing ∼90% depletion in the cortex.

**Fig. S2. qPCR validation.** (A) qPCR Cq values compared to the corresponding log2 of the RNAseq read counts. Each datapoint represents the mean +/- SEM for a miRNA-experimental condition combination. qPCR value was taken as the average of n = 3 technical replicates for 54 unique miRNA-sample combinations. qPCR values >40 were excluded from analysis. Correlation evaluated using Pearson’s r. (B) miRNAs evaluated using qPCR to compare against sequencing.

**Fig. S3. Extended single nuclei characterization.** (A) Dot plot showing the top 3 marker genes for differentiating nuclei types. Note that these marker genes are representative and were not explicitly used to define nuclei clusters; CellTypist uses a holistic logistic regression-based approach. (B) Distribution of nuclei types across experimental conditions. (C) *Gria1* and *Gria2* (which encode GluA1 and GluA2, respectively) expression as a function of experimental condition across nuclei types.

**Fig. S4. Volcano plots for within-Vehicle comparisons.** S: Sham; C: Concussion; PC: Preconditioned Concussion.

**Fig. S5. Volcano plots for within-PLX5622 comparisons.** S: Sham; C: Concussion; PC: Preconditioned Concussion.

**Fig. S6. Volcano plots for PLX5622 vs Vehicle.** S: Sham; C: Concussion; PC: Preconditioned Concussion.

**Fig. S7. Cytokine panel shows no significant injury or PLX5622-mediated differences.** Serum cytokine levels at 9 days post-injury for Vehicle and PLX5622 mice across sham, concussion, and preconditioned concussion. Mixed effects model with Geisser-Greenhouse correction and Tukey’s multiple comparisons test. Data are presented as mean +/- SEM. * p<0.05.

**Fig. S8. PLX5622 treatment decreases ambulation and impairs fear memory independently of injury.** (A) PLX5622-treated mice exhibit decreased ambulation in an open field regardless of injury condition. (B) There is no difference in proportional regional exploration in an open field due to injury or PLX5622 treatment. (C) PLX5622-treated mice exhibit decreased freeze percentage regardless of injury condition. Two-way ANOVA with Sidak’s test for multiple comparisons. Data are presented as mean +/-SEM. * p<0.05, ** p<0.01.

**TableS1_Injury_miRNAs.xlsx**

**Table S1.** Complete list of GO: BP terms associated with differential miRNA expression for injury biomarkers.

**TableS2_InjuryHistory_miRNAs.xlsx**

**Table S2.** Complete list of GO: BP terms associated with differential miRNA expression for injury history biomarkers.

**TableS3_PLXvsVeh_miRNAs.xlsx**

**Table S3.** Complete list of GO: BP terms associated with differential miRNA expression for concussed mice with PLX5622 vs Vehicle diet.

**TableS4_snRNAseq_DEGs.xlsx**

**Table S4.** Complete list of differentially expressed genes (FDR-adjusted p < 0.1), including log2FC, FDR-adjusted p-value, average relative expression, and percent expression.

**TableS5_NOR_miRNAs.xlsx**

**Table S5.** Complete list of GO: BP terms associated with LASSO panel of 13 novel object recognition (NOR)-associated miRNAs.

