## Additional Files for "Microglia modulate concussion biomarkers and cognitive recovery in male mice"

### Fig. S**1. Validation of microglial depletion.** Representative coronal sections at -1.5-2 mm from bregma following 10d of (A) Vehicle or (B) PLX5622 treatment showing ~90% depletion in the cortex.

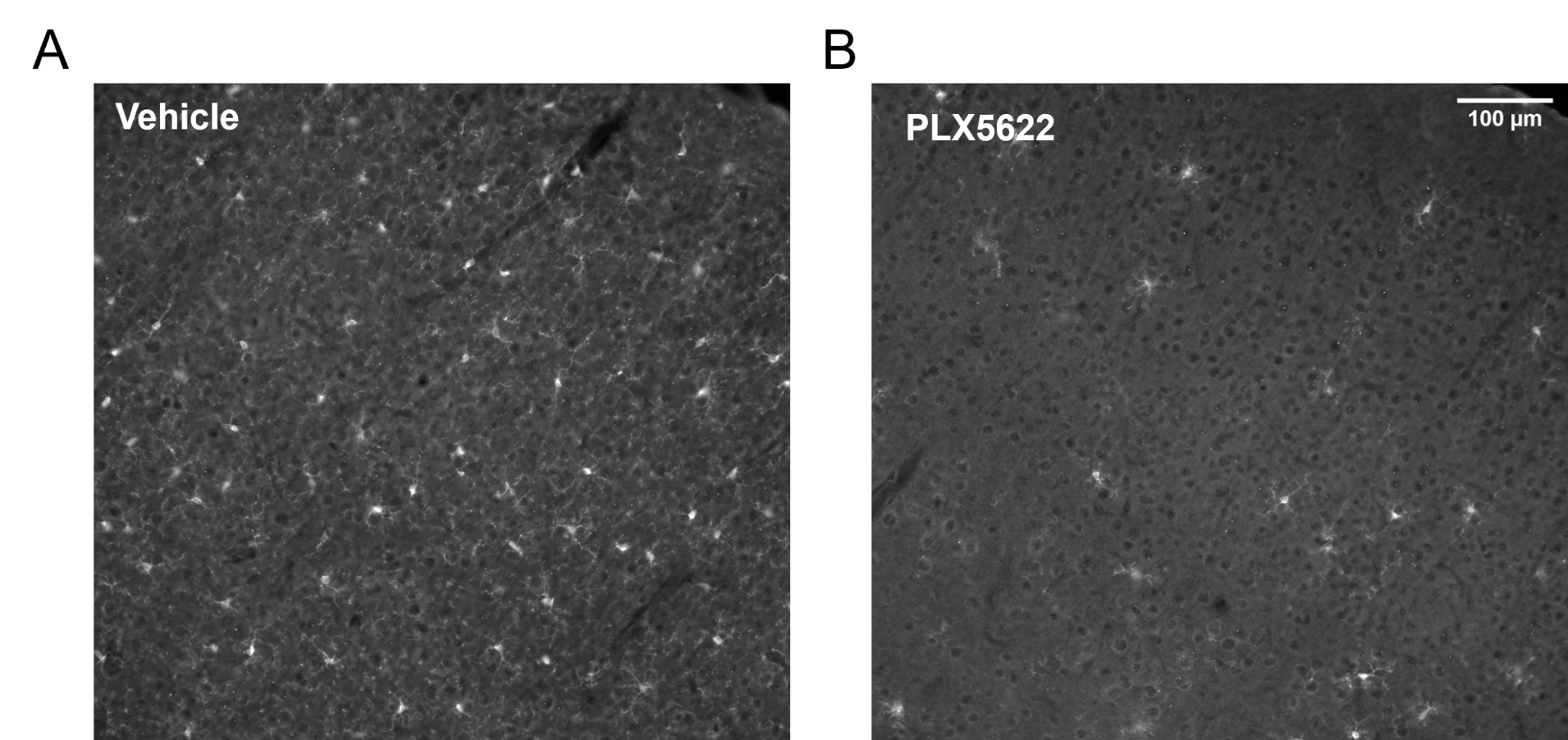

### Fig. S2. qPCR validation. (A) qPCR Cq values compared to the corresponding log2 of the RNAseq read counts. Each datapoint represents the mean +/- SEM for a miRNA-experimental condition combination. qPCR value was taken as the average of n = 3 technical replicates for 54 unique miRNA-sample combinations. qPCR values >40 were excluded from analysis. Correlation evaluated using Pearson’s r. (B) miRNAs evaluated using qPCR to compare against sequencing.

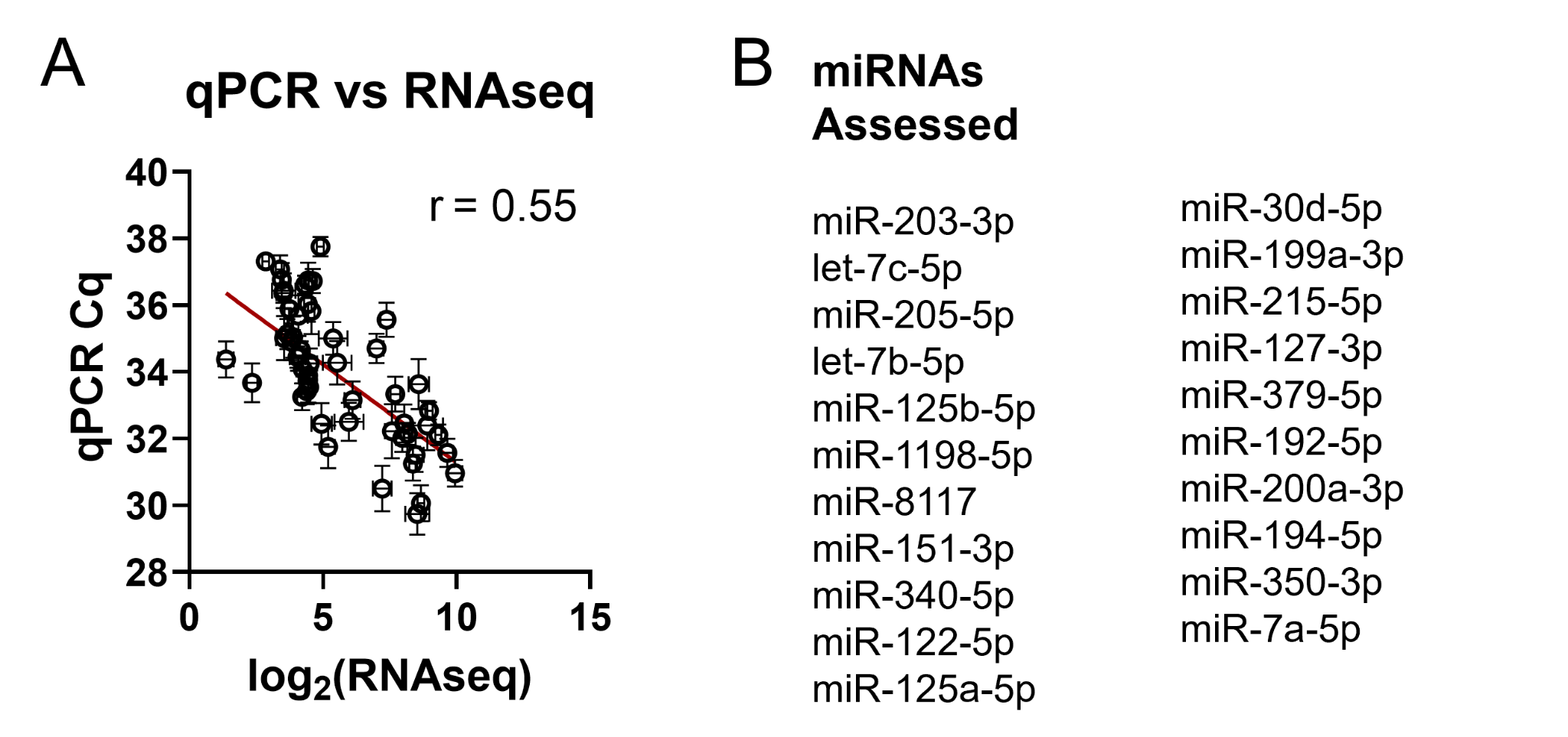

### Fig. S3. Extended single nuclei characterization. (A) Dot plot showing the top 3 marker genes for differentiating nuclei types. Note that these marker genes are representative and were not explicitly used to define nuclei clusters; CellTypist uses a holistic logistic regression-based approach. (B) Distribution and counts of nuclei types across experimental conditions. (C) *Gria1* and *Gria2* (which encode GluA1 and GluA2, respectively) expression as a function of experimental condition across nuclei types.

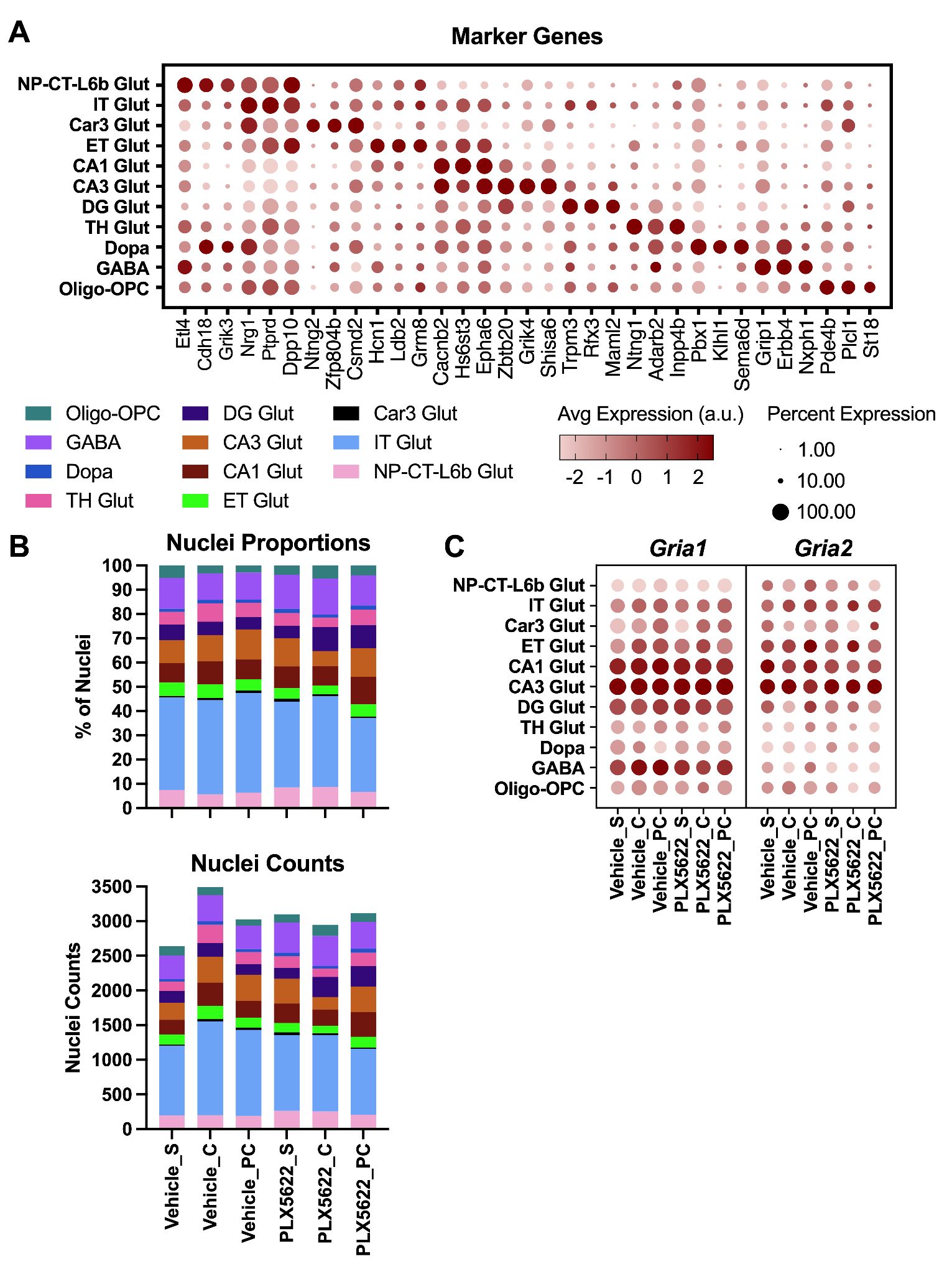

### Fig. S4. Volcano plots for within-Vehicle comparisons. S: Sham; C: Concussion; PC: Preconditioned Concussion.

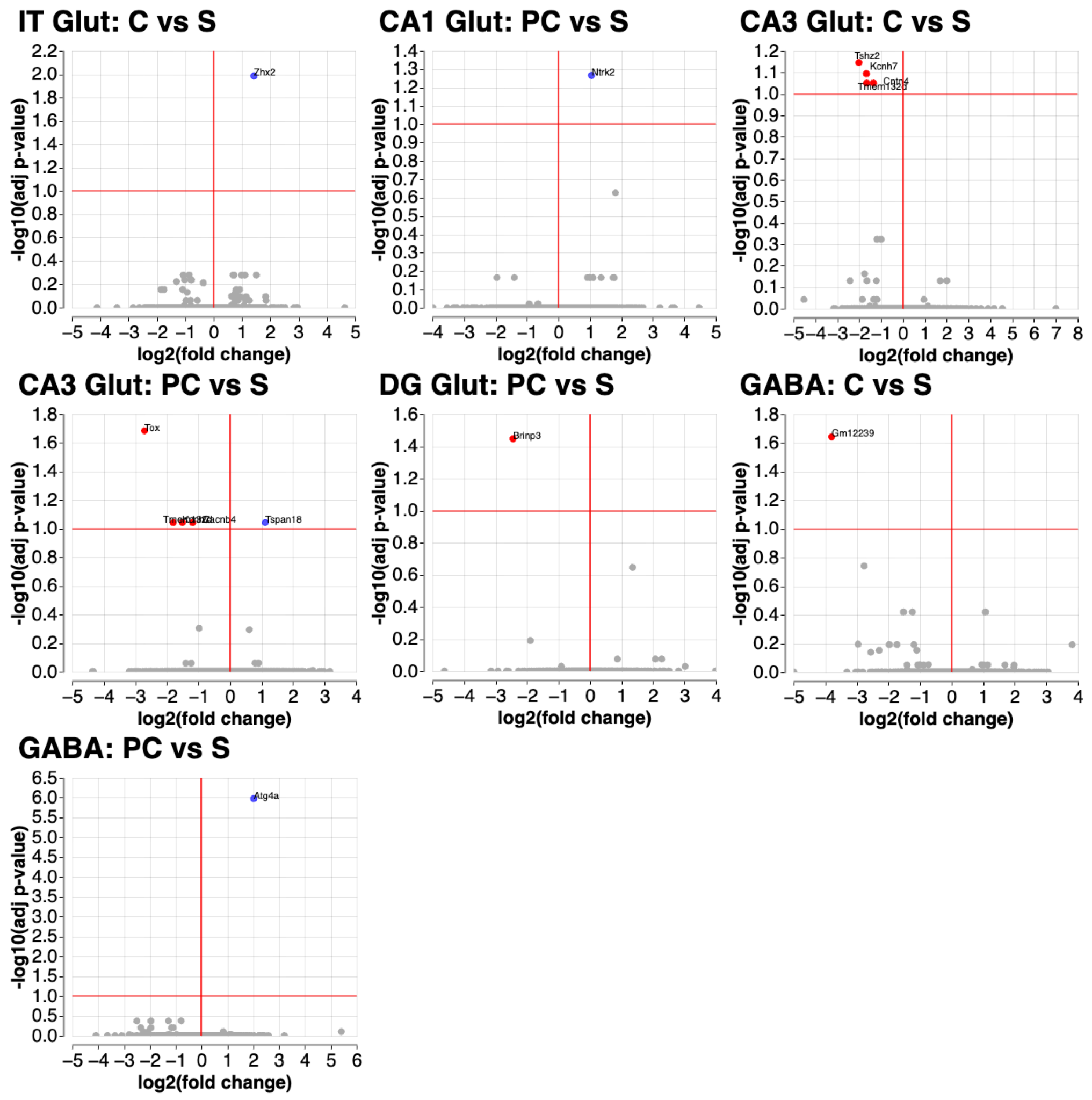

### Fig. S5. Volcano plots for within-PLX5622 comparisons. S: Sham; C: Concussion; PC: Preconditioned Concussion.

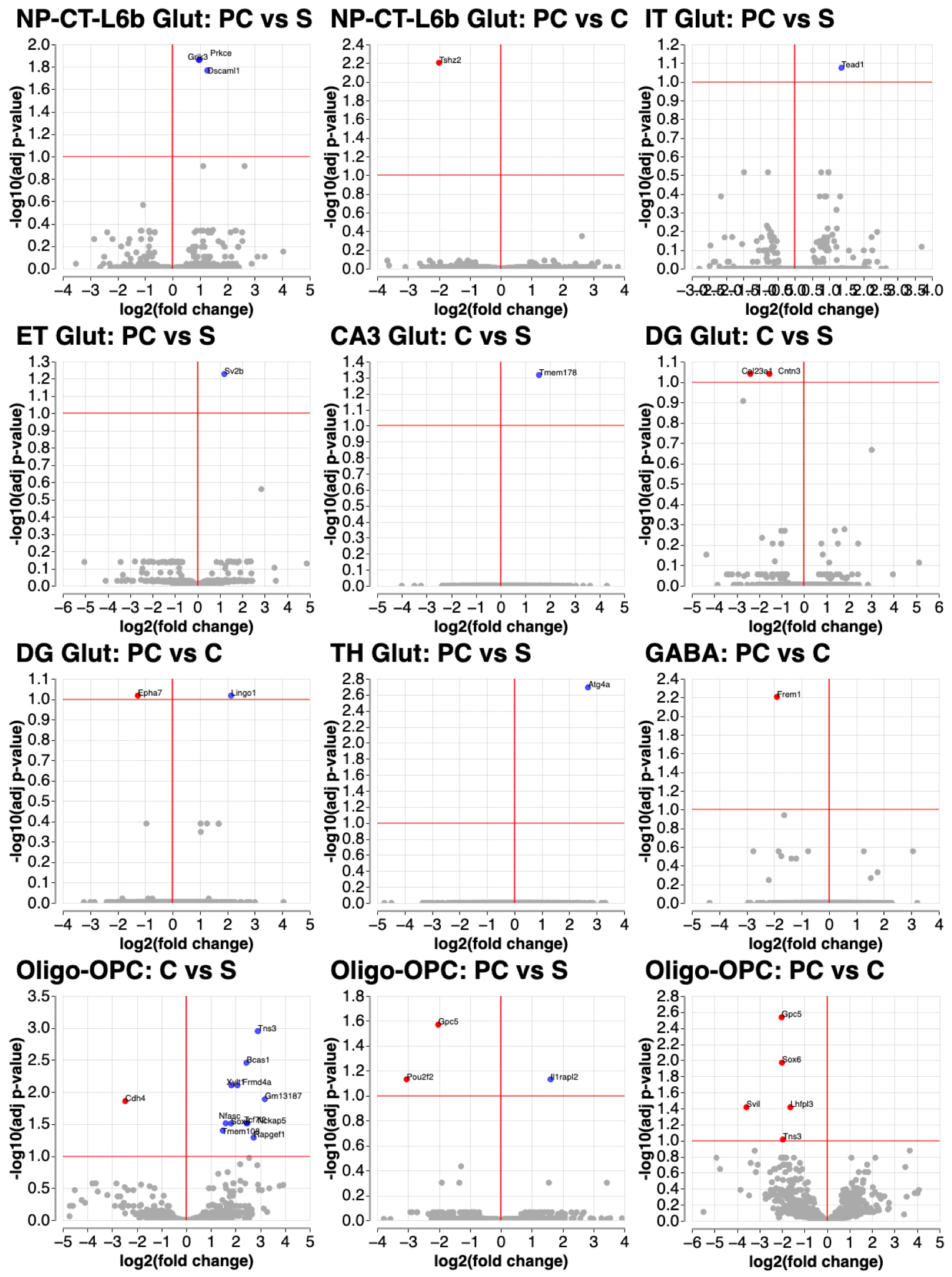

### Fig. S6. Volcano plots for PLX5622 vs Vehicle. S: Sham; C: Concussion; PC: Preconditioned Concussion.

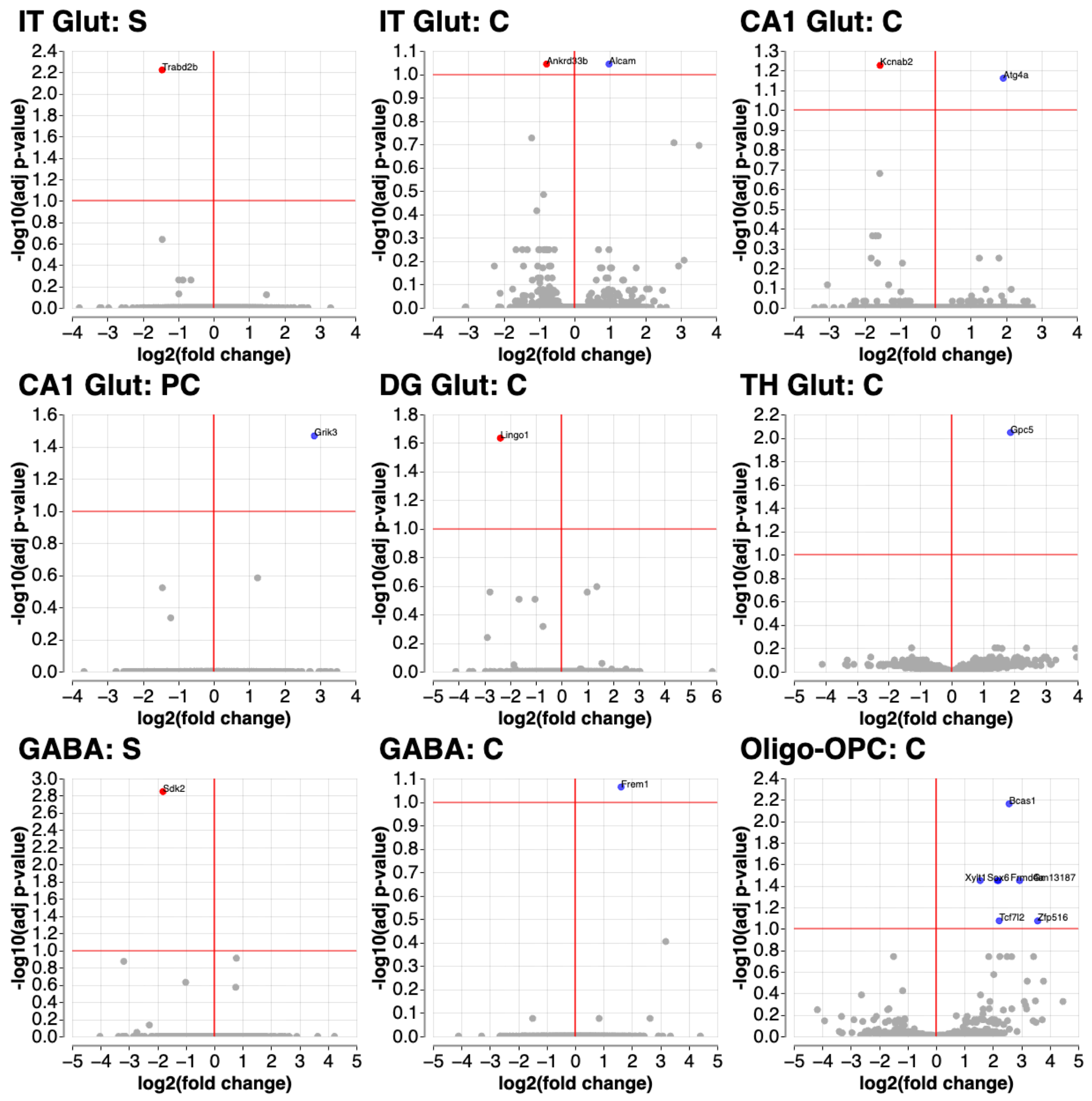

### Fig. S7**. Cytokine panel shows no significant injury or PLX5622-mediated differences.** Serum cytokine levels at 9 days post-injury for Vehicle and PLX5622 mice across sham, concussion, and preconditioned concussion. Mixed effects model with Geisser-Greenhouse correction and Tukey’s multiple comparisons test. Data are presented as mean +/- SEM. * p<0.05.

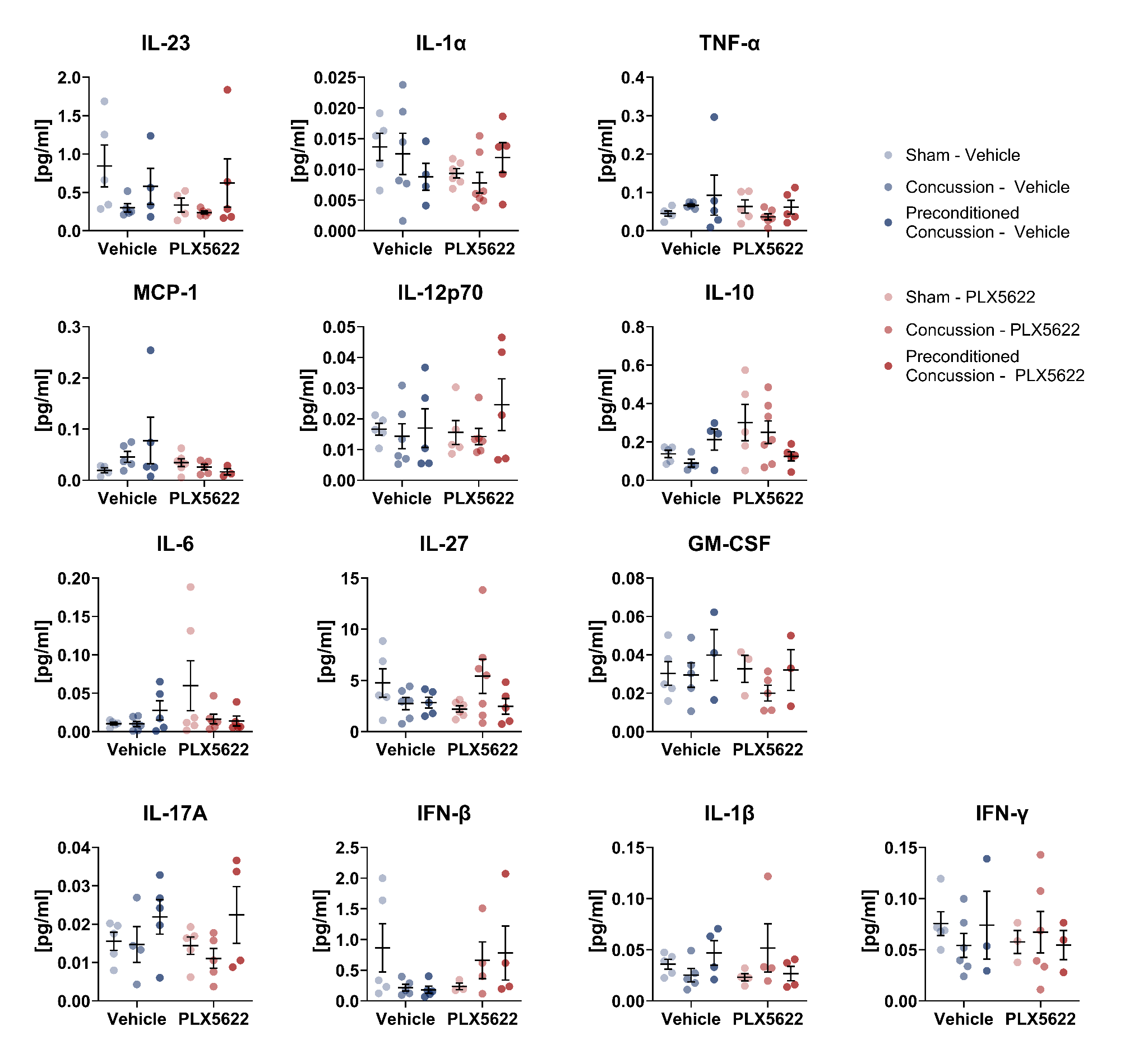

### Fig. S8. PLX5622 treatment decreases ambulation and impairs fear memory independently of injury. (A) PLX5622-treated mice exhibit decreased ambulation in an open field regardless of injury condition. (B) There is no difference in proportional regional exploration in an open field due to injury or PLX5622 treatment. (C) PLX5622-treated mice exhibit decreased freeze percentage regardless of injury condition. Two-way ANOVA with Sidak’s test for multiple comparisons. Data are presented as mean +/- SEM. * p<0.05, ** p<0.01.

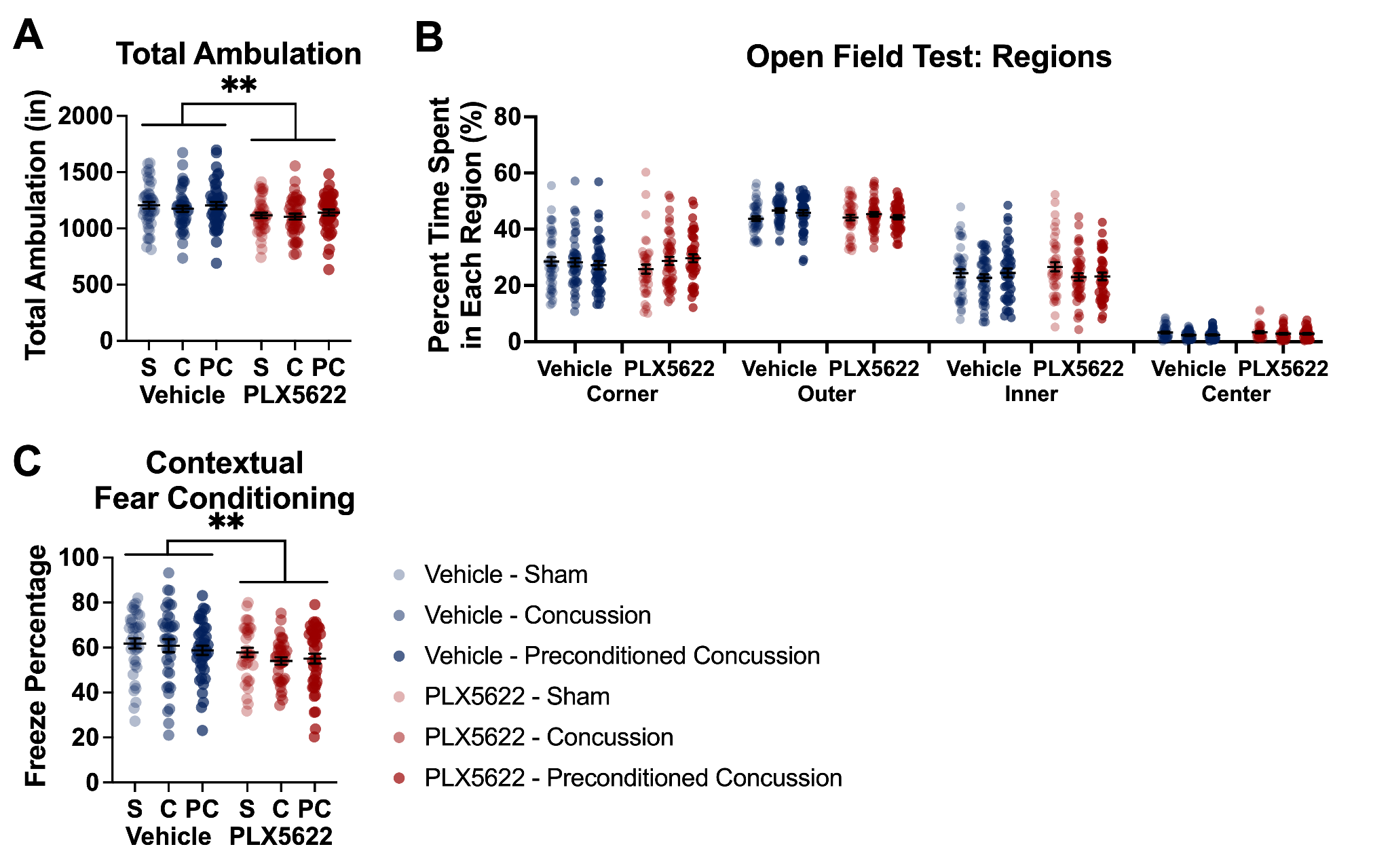
